# Fine-tuning of coumaric acid synthesis to increase naringenin production in yeast

**DOI:** 10.1101/2022.06.20.496858

**Authors:** Jiwei Mao, Marta Tous Mohedano, Xiaowei Li, Quanli Liu, Jens Nielsen, Verena Siewers, Yun Chen

**Affiliations:** Department of Biology and Biological Engineering, Chalmers University of Technology, SE412 96 Gothenburg, Sweden; BioInnovation Institute, DK2200 Copenhagen N, Denmark

**Keywords:** Synthetic biology, Metabolic engineering, Dynamic control, Intermediate transport, Natural products

## Abstract

(2*S*)-Naringenin is a key precursor for biosynthesis of various high-value flavonoids and possesses a variety of nutritional and pharmaceutical properties on human health. Systematic optimization approaches have been employed to improve (2*S*)-naringenin production in different microbial hosts. However, very few studies have focused on the spatiotemporal distribution of (2*S*)-naringenin and related pathway intermediate *p*-coumaric acid, which is an important factor for efficient production. Here, we show that fine-turning of *p*-coumaric acid synthesis enables alleviated cell burden and improved (2*S*)-naringenin production in yeast. First, we systematically optimized the (2*S*)-naringenin biosynthetic pathway by alleviating the bottleneck downstream of *p*-coumaric acid and increasing malonyl-CoA supply, which improved (2*S*)-naringenin production but significant amounts of *p*-coumaric acid still accumulated outside the cell. We further established a dual dynamic control system through combing a malonyl-CoA biosensor regulator and an RNAi strategy, to autonomously control the synthesis of *p*-coumaric acid and downregulate a pathway competing for malonyl-CoA. The optimized strains remarkably decreased extracellular accumulation of *p*-coumaric acid and simultaneously improved (2*S*)- naringenin production. Finally, production of 933 mg/L of (2*S*)-naringenin could be achieved by using minimal medium with negligible accumulation of *p*-coumaric acid. Our work highlights the importance of systematic control of pathway intermediates for efficient microbial production of plant natural products.

## Introduction

Flavonoids are the largest classes of polyphenolic secondary metabolites in plants, representing an important resource of therapeutics and nutraceuticals due to their antioxidant, anti-inflammatory, antitumor activities, liver-protective properties as well as memory enhancement (1, 2). The (2*S*)- naringenin is one of the most important scaffolds for producing a wide variety of subgroups of flavonoids. The formation of (2*S*)-naringenin begins with the conversion of L-tyrosine or L-phenylalanine to *p*-coumaric acid through the action of tyrosine ammonia lyase (TAL) or phenylalanine ammonia lyase (PAL) combined with cinnamic acid hydroxylase (C4H), respectively. *p*-Coumaric acid is subsequently converted by *p*-coumaroyl-CoA ligase (4CL) to form the 4-coumaroyl-CoA. Next, one molecule of 4-coumaroyl-CoA is condensed with three molecules of malonyl-CoA to generate naringenin chalcone by the type III polyketide synthase, chalcone synthase (CHS). In the final step, the naringenin chalcone is converted to (2*S*)-naringenin by the action of chalcone isomerase (CHI) (Fig. 1A).

**Figure 1.**
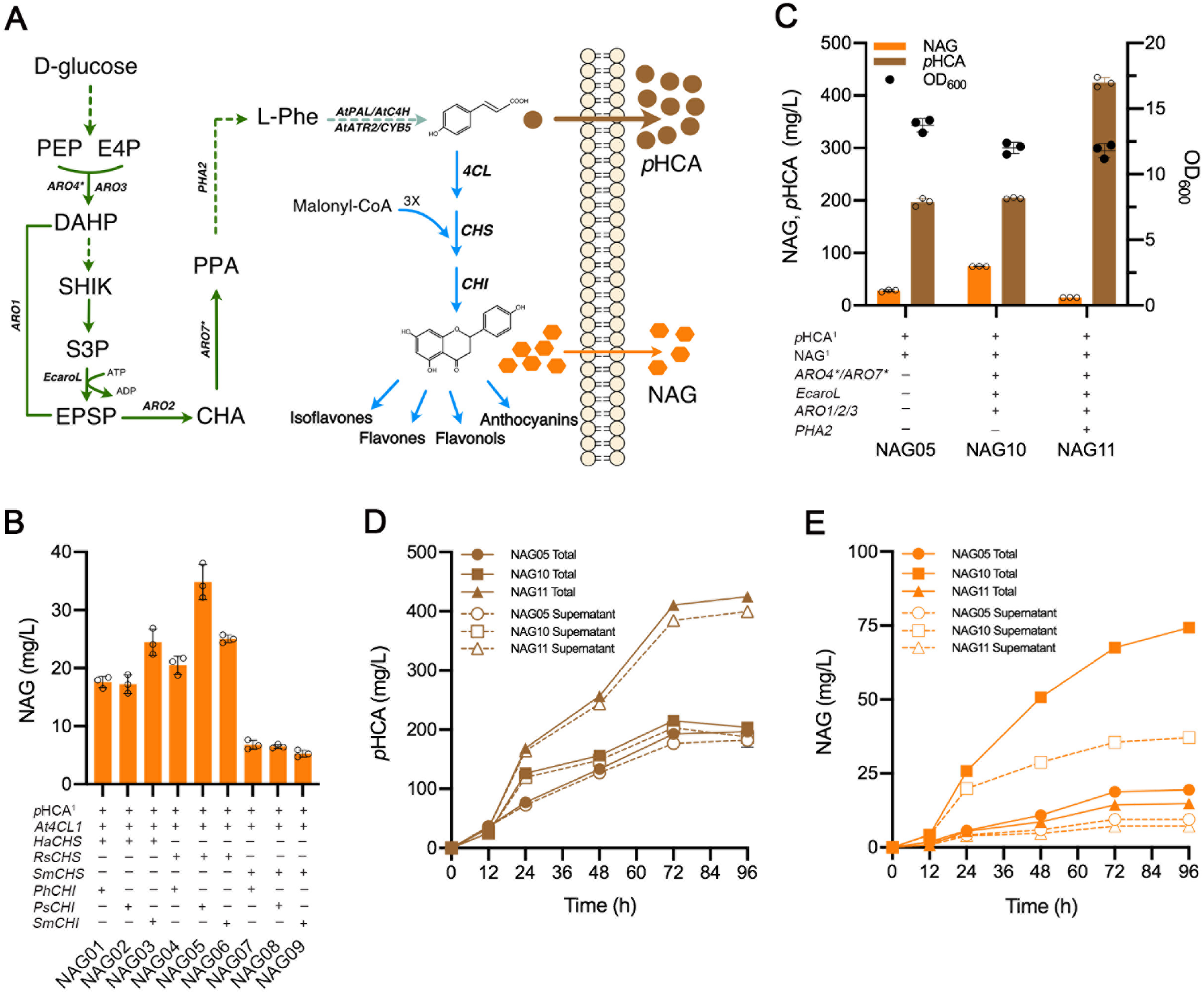
Spatial mismatch of *p*-coumaric acid with (2*S*)-naringenin biosynthesis. (A) Schematic diagram of the biosynthetic pathway leading to the production of (2*S*)-naringenin in *S. cerevisiae*. Solid lines represent a single step; dotted lines indicate multiple steps; the blue arrows represent the core (2*S*)-naringenin biosynthetic pathway and a variety of plant specific flavonoids derived from (2*S*)- naringenin. *ARO3*, DAHP synthase; *ARO4\**, L-tyrosine-feedback-insensitive DAHP synthase (*ARO4*^*K229L*^); *ARO1*, pentafunctional aromatic protein; *EcoroL*, shikimate kinase from *E. coli*; *ARO2*, chorismate synthase; *ARO7\**, L-tyrosine-feedback insensitive chorismate mutase (*ARO7*^*G141S*^); *AtPAL2*, phenylalanine ammonia lyase, *AtC4H*, cinnamic acid hydroxylase and *AtATR2*, cytochrome P450 reductase, from *Arabidopsis thaliana*; *CYB5*, yeast native cytochrome b5; *4CL, p*-coumaroyl-CoA ligase; *CHS*, chalcone synthase; *CHI*, chalcone isomerase. E4P, erythrose-4-phosphate; PEP, phosphoenolpyruvate; DAHP, 3-deoxy-D-arabino-2-heptulosonic acid 7-phosphate; SHIK, shikimate; S3P, shikimate-3-phosphate; EPSP, 5-enolpyruvyl-shikimate-3-phosphate; CHA, chorismic acid; PPA, prephenate; L-Phe, L-phenylalanine; *p*HCA, *p*-coumaric acid; NAG, (2*S*)-naringenin. (B) (2*S*)- naringenin production by engineered strain expressing the core (2*S*)-naringenin biosynthetic genes from different sources. *p*HCA^1^ represents *p*-coumaric acid biosynthesis pathway consisting of *AtPAL2, AtC4H, AtATR2* and *CYB5*. (C) Metabolic engineering to enhance the metabolic flux from glucose to *p*-coumaric acid. NAG^1^ represents the basic core (2*S*)-naringenin biosynthetic pathway, consisting of one copy of *At4CL1* from *A. thaliana, RsCHS* from *Rhododendron simsii* and *PsCHI* from *Paeonia suffruticosa*. (D)(E) Intracellular and extracellular content of *p*-coumaric acid and (2*S*)-naringenin in different engineered strains. The cells were cultivated in defined minimal medium with 30 g/L glucose as the sole carbon source, and cultures were sampled after 96 h of growth for *p*-coumaric acid and (2*S*)- naringenin detection. All data represent the mean of n = 3 biologically independent samples and error bars show standard deviation.

For robust production of (2*S*)-naringenin, its biosynthetic pathway has been reconstituted into different microorganisms including *Escherichia coli* (3, 4), *Corynebacterium glutamicum (5), Streptomyces venezuelae* (6), *Yarrowia lipolytica* (7, 8) and *Saccharomyces cerevisiae* (9-12). Progress has been made to improve (2*S*)-naringenin production by supplementation of precursor amino acids (L-tyrosine, L-phenylalanine) or *p*-coumaric acid, or by *de novo* synthesis from simple carbon source like glucose (13-16). While most efforts have been centered on optimizing the (2*S*)-naringenin biosynthetic pathway, very few studies have focused on the spatiotemporal distribution of its intermediate such as *p*-coumaric acid and the product (2*S*)-naringenin, an important aspect for efficient production.

In plants, the biosynthesis, storage, and function of secondary metabolites to fulfill their biological roles often involve different cellular localizations and processes. The subcellular translocation of secondary metabolites is likely to occur by active transport processes, but the specific transporters involved have not been satisfactorily identified (17, 18). Furthermore, when transferring plant pathway enzymes into microbial host the cellular environment can be different, which may lead to distinct enzyme activities and intermediate transportation processes. For instance, when plant alkaloid pathways distributed across different organelles and tissues were reconstituted in yeast, it showed the production of previously inaccessible intermediates due to unicellular characteristic of yeast (19), or limitations in cellular transport of intermediates as lacking of appropriate transporters in host cell (20). On the other hand, plant pathway intermediates may be exported out of host cells, which will result in lower intracellular concentrations, and can be deleterious to the metabolic flux through the pathway especially for downstream of slower enzymatic steps. In addition, the efficient export of pathway intermediates can lead to carbon loss and accumulation of toxic intermediates, which are important obstacles hinder efficient microbial production.

In this study, we quantified the intracellular and extracellular content of (2*S*)-naringenin and the intermediate *p*-coumaric acid and found that more than 90% of the *p*-coumaric acid produced was exported outside the cell at all production levels. To overcome the issue with *p*-coumaric acid secretion, we first optimized the (2*S*)-naringenin biosynthetic pathway by alleviating the bottleneck downstream of *p*-coumaric acid and increasing the supply of malonyl-CoA, which resulted in improved (2*S*)-naringenin production, but significant accumulation of *p*-coumaric acid still existed. To further address this, we employed a dual dynamic control system through a malonyl-CoA biosensor in combination with an RNAi strategy to autonomously balance the supply of malonyl-CoA and *p*-coumaric acid. This significantly reduced the accumulation of *p*-coumaric acid and increased the titer of (2*S*)-naringenin in the resultant strain. Ultimately, a high-level production of (2*S*)-naringenin was achieved, up to 933 mg/L under fed-batch fermentation conditions. We believe that the strategies developed could be potentially applicable for other microbial cell factories and natural products of interest. Also, our work demonstrates the importance of controlling pathway intermediates synthesis for microbial based production of plant natural products.

## Results

### Spatial mismatch of *p*-coumaric acid with (2*S*)-naringenin production

As shown in Fig. 1A, a L-phenylalanine-based pathway was used for *de novo* synthesis of *p*-coumaric acid as described earlier (21). To further construct the (2*S*)-naringenin biosynthetic pathway, we screened different CHSs and CHIs from different sources (Fig. 1B). Among different combinations of CHSs and CHIs, a strain containing *RsCHS* from *Rhododendron simsii* and *PsCHI* from *Paeonia suffruticosa* was found to be best resulting in production of 35 mg/l of (2*S*)-naringenin. As *p*-coumaric acid is an essential precursor for (2*S*)-naringenin production, we next expressed feedback resistant variants of *ARO4\** and *ARO7\**, and additional copies of *ARO1/2/3* together with *Ec*aroL from *E. coli*, which has been previously shown to significantly improve *p*-coumaric acid production (21). This resulted in increased (2*S*)-naringenin production, up to 68 mg/L in strain NAG10 (Fig. 1C). However, further enhancement of *p*-coumaric acid production by additionally expressing *PHA2* led to decreased (2*S*)- naringenin production in NAG11, while at the same time *p*-coumaric acid production increased significantly, up to 350 mg/L (Fig. 1C).

When we examined the whole production process it was found that more than 90% of the *p*-coumaric acid produced was secreted out of the cell independent of production level (Fig. 1D), indicating that yeast cells are very efficient in exporting *p*-coumaric acid. On the other hand, (2*S*)-naringenin seemed to be secreted mainly during the first 24 hours and then started to accumulate intracellularly (Fig. 1E). Moreover, the accumulation of *p*-coumaric acid didn’t show a significant reduction trend even during prolonged cultivation. This indicated that extracellular *p*-coumaric acid was not efficiently re-imported by the cells for (2*S*)-naringenin biosynthesis. Comparing strain NAG05, NAG10 and NAG11, which have different capabilities of *p*-coumaric acid production, only the intermediate level of *p*-coumaric acid production (NAG10) improved (2*S*)-naringenin production. Nevertheless, still excessive *p*-coumaric acid (204 mg/L) was secreted by the NAG10 strain. This could imply that the pathway downstream of *p*-coumaric acid is insufficient and/or the other precursor malonyl-CoA is limiting.

### Alleviating the bottleneck downstream of *p*-coumaric acid improves (2*S*)-naringenin production

We firstly checked if the expression level of downstream enzymes of *p*-coumaric acid is limiting for (2*S*)- naringenin production. Variant gene copies of *4CL, CHS* and *CHI* were inserted into the genome of the NAG10 strain (Fig. 2A). Overexpressing all three enzymes individually increased (2*S*)-naringenin production, while simultaneous overexpression of *CHS* and *CHI* had a more pronounced effect. We thus grouped *CHS* and *CHI* together and systematically evaluated the effect of overexpression of (*4CL*: *CHS&CHI*) in the range of 1 to 4 copies (Fig. 2B). Generally, with a fixed *4CL* level increasing the expression of *CHS&CHI* from 1 to 4 copies, (2*S*)-naringenin production increased accordingly, strongly suggesting that these two enzymes are not sufficient. When *4CL* increased from 3 to 4 copies, the trend of increasing (2*S*)-naringenin production seemed to be saturated, and further increasing the gene copy number of either *4CL* or *CHS&CHI* did not improve (2*S*)-naringenin production anymore (Fig. S1). Eventually, the optimal ratio of *4CL* and *CHS&CHI* in NAG3-4 was 3:4, which led to production of 209 mg/L of (2*S*)-naringenin, a 3-fold improvement relative to the parental strain NAG10.

**Figure 2.**
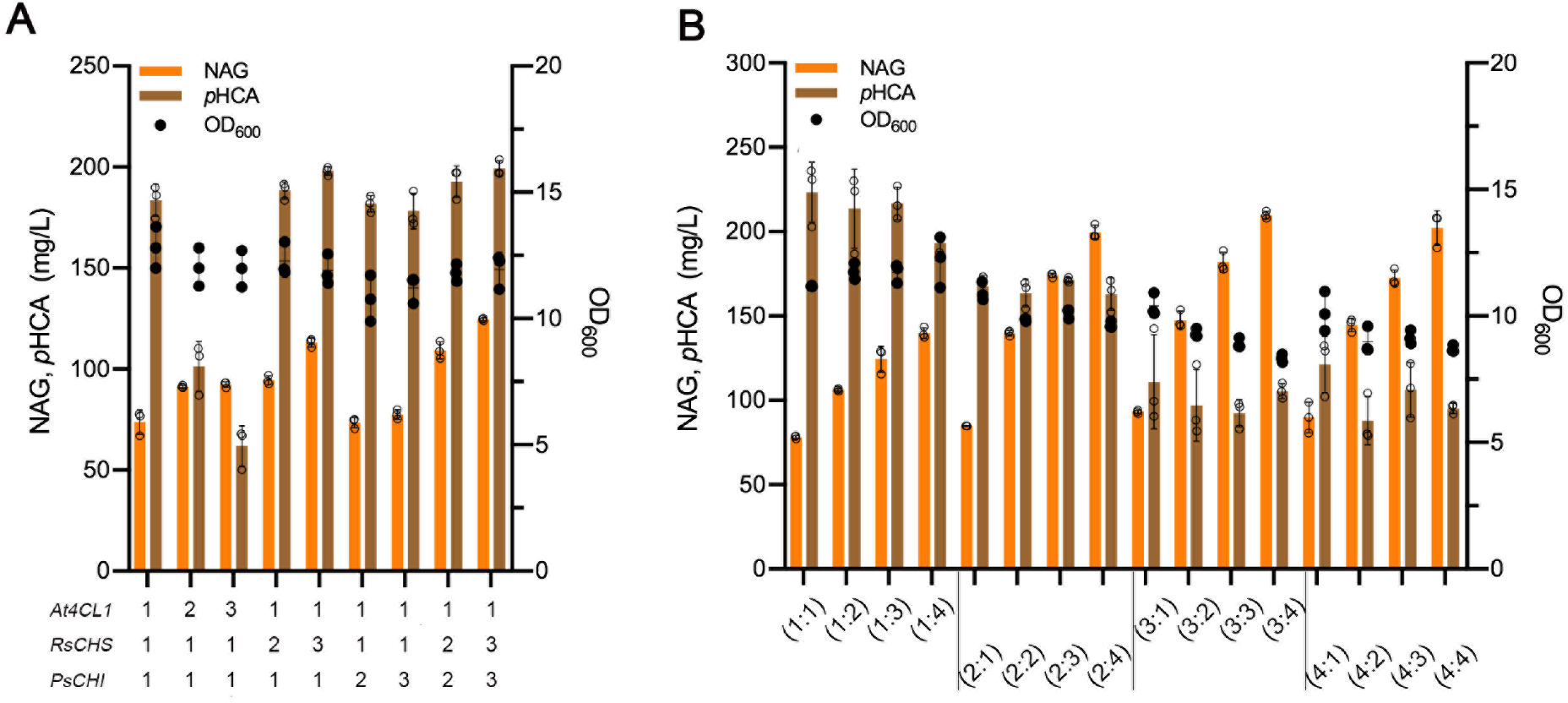
Optimization of the core (2*S*)-naringenin biosynthetic pathway. (A) Rate-limiting steps were analyzed by gradually increasing the gene copy number of each step. (B) Fine-tuning the expression of 4CL and CHS&CHI to improve (2*S*)-naringenin production. The numbers (n : n) indicate the respective gene copy numbers of *4CL* and *CHS&CHI*. Cells were grown in a defined minimal medium with 30 g/L glucose as the sole carbon source, and cultures were sampled after 96 h of growth for metabolite analysis. All data represent the mean of n = 3 biologically independent samples and error bars show standard deviation.

Interestingly, when the copy number of *4CL* increased from 1 to 3 there was also a trend towards decreased *p*-coumaric acid accumulation (Fig. 2B), whereas its accumulation was not significantly affected by the expression level of *CHS&CHI*. This could be due to the competition between a potential transporter and 4CL for the same substrate of *p*-coumaric acid; if it is not captured by 4CL then it will be secreted out of the cell. Nonetheless, when *4CL* increased from 3 to 4 copies, there was no further decrease of *p*-coumaric acid accumulation. These results indicated that the accumulation of *p*-coumaric acid cannot be eliminated completely by expanding the expression level of downstream enzymes.

### Elevated malonyl-CoA supply increases (2*S*)-naringenin production

Malonyl-CoA is another important precursor for flavonoid biosynthesis, and enhancing malonyl-CoA flux may thus push the biosynthesis pathway to improve (2*S*)-naringenin production while reducing *p*-coumaric acid accumulation. To evaluate the effect of malonyl-CoA supply on (2*S*)-naringenin production, we used three approaches, i) a mutant version of Acc1(catalyzing the formation of malonyl-CoA from acetyl-CoA), mAcc1^*^ (Acc1^ser659ala,ser1157ala^) with enhanced activity (22); ii) an alternative pathway from *Rhizobium trifolii* containing a malonate transporter (matC) and activation (matB) for malonyl-CoA formation (23); and iii) a combination of these two methods. We first tested these strategies in the background of NAG10 strain (Fig. 3A). Expression of either mAcc1^*^ or matBC improved (2*S*)- naringenin production, while the combination of these two strategies gave the highest production at 122 mg/L of (2*S*)-naringenin, representing an 80% increase compared with NAG10. However, the increase was less efficient than the enhanced *p*-coumaric acid downstream pathway, which resulted in 209 mg/L of (2*S*)-naringenin, pointing that the efficiency of the *p*-coumaric acid downstream pathway is more severely limited. Surprisingly, (2*S*)-naringenin production was not significantly increased further when we tested these malonyl-CoA supply strategies in the background of NAG3-4 strain (Fig. 3B). However, the accumulation of *p*-coumaric acid was further increased when expressing mAcc1^*^ and matBC in both background of NAG10 and NAG3-4 strains (Fig. 3A, 3B). At the same time, biomass was significantly decreased especially in the strain with both mAcc1^*^ and matBC, which may indicate an imbalance in the supply of precursor *p*-coumaric acid and malonyl-CoA.

**Figure 3.**
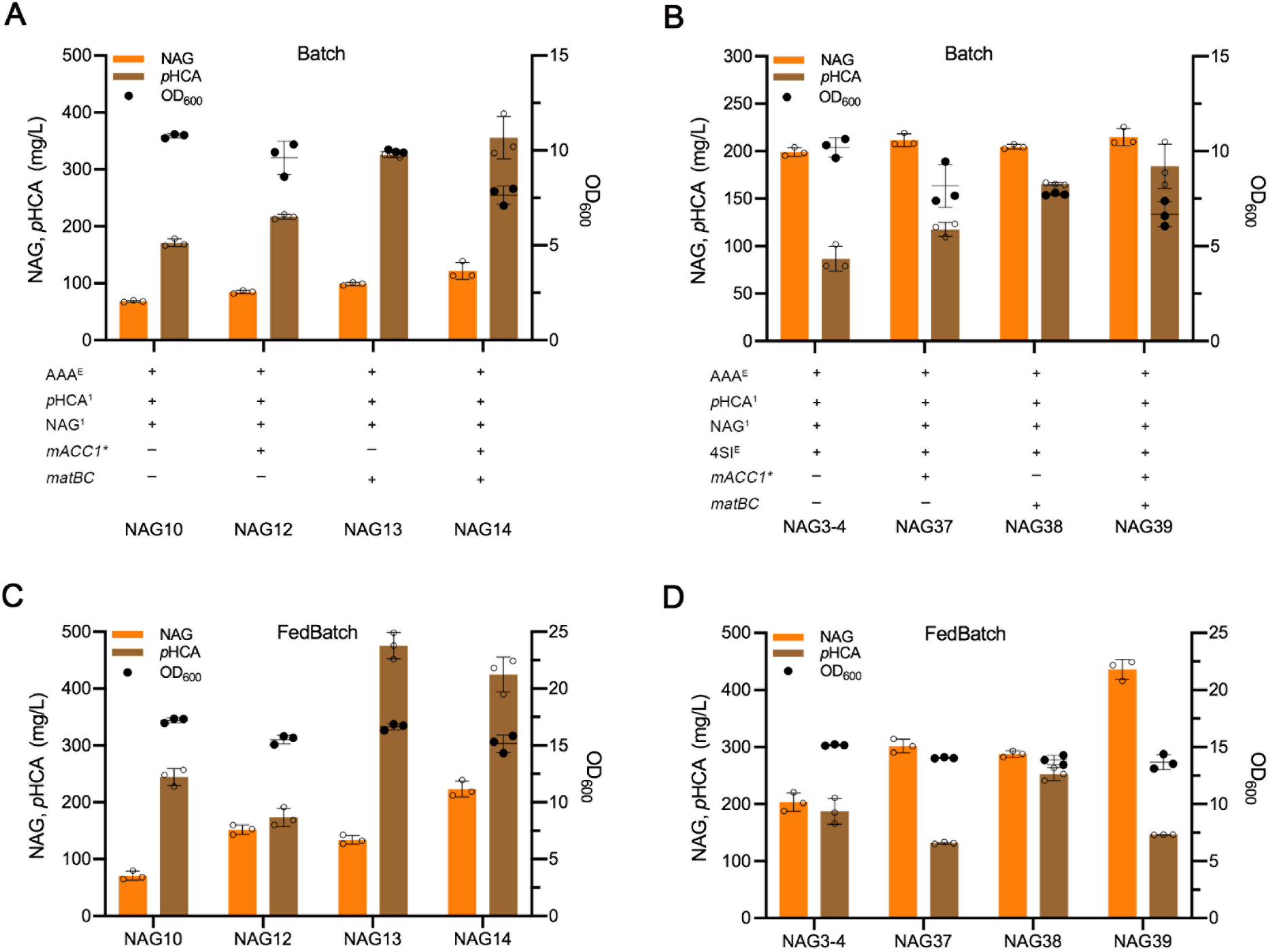
Improving the malonyl-CoA availability to increase(2S)-naringenin production. (A)(B) Production of (2*S*)-naringenin in batch condition by strains with different strategies to enhance the supply of malonyl-CoA. AAA^E^ represents the overexpression of L-phenylalanine biosynthesis pathway. *p*HCA^1^ refers to *p*-coumaric acid biosynthesis pathway. NAG^1^ represents the basic core (2*S*)-naringenin biosynthetic pathway, consisting of one copy of *At4CL1* from *A. thaliana, RsCHS* from *Rhododendron simsii* and *PsCHI* from *Paeonia suffruticosa*. 4SI^E^ represents the optimization of the core (2*S*)-naringenin biosynthetic pathway, including two copies of *At4CL1*, three copies of *RsCHS* and *PsCHI. mACC1\** represents the overexpression of *ACC1*^*S659A, S1157A*^. matB,C represents malonate assimilation pathway. Cells were grown in defined minimal medium with 30 g/L glucose, and with 2 g/L sodium malonate dibasic supplementation when required. Cultures were sampled after 96 h of cultivation for metabolite detection. (C)(D) Process optimization for (2*S*)-naringenin production by malonyl-CoA enhancing strains derived from NAG10 and NAG3-4 background strains, respectively. Cells were grown in defined minimal medium with six tablets of FeedBeads as the sole carbon source, and with 2 g/L sodium malonate dibasic supplementation when required. Cultures were sampled after 96 h of cultivation for metabolite analysis. All data represent the mean of n= 3 biologically independent samples and error bars show standard deviation.

We further examined these strains in a fed-batch-like condition with slow release of glucose, here, a more substantial increase was found in both background of NAG10 and NAG3-4 strains (Fig. 3C, 3D). Expression of either mAcc1^*^ or matBC improved (2*S*)-naringenin production by about 2-fold, while the combination of the two increased (2*S*)-naringenin production by more than 3-fold, up to 223 mg/L (2*S*)- naringenin in strain NAG14. Although in batch conditions there was no increase in NAG3-4 background strains, a significant increase was observed under fed-batch mimicking conditions. The combination of mAcc1^*^and matBC resulted in the highest production up to 436 mg/L in NAG39, representing a 2-fold improvement relative to the NAG3-4 background strain. However, this increase was less than that of NAG10 background strain, suggesting that malonyl-CoA still was the limiting factor in NAG39, in which the downstream capability of *p*-coumaric acid was increased. Compared with the normal batch conditions, the fed-batch like cultivation limits the overflow of metabolism through continuously releasing glucose at a low level. The improved (2*S*)-naringenin production may be attributed to the *in vivo* balanced flux between *p*-coumaric acid production and malonyl-CoA supply under glucose-limited cultivation. However, under this condition, *p*-coumaric acid was still accumulated and the biomass was significantly reduced in the NAG39 strain.

### Dynamic regulation of *p*-coumaric acid synthesis reduces its accumulation

To autonomously balance the metabolic flux between *p*-coumaric acid production and malonyl-CoA supply, we implemented a malonyl-CoA biosensor to dynamically control the production rate of *p*-coumaric acid during cell growth (Fig. 4A). The FapR−*fapO* system from *Bacillus subtilis* (24), has been successfully applied in mammalian cells (25), *E. coli* (26) and *S. cerevisiae* (27) to build the malonyl-CoA responsive sensors. Additionally, in our previous studies, a set of synthetic promoters were developed with varied sensitivity and dynamic ranges in response to malonyl-CoA by changing number and/or inserting position of *fapO* in native yeast promoters (28). Thus, these modified promoters were chosen to control the expression of *AtPAL*, which is the first enzyme of the pathway leading to *p*-coumaric acid production.

**Figure 4.**
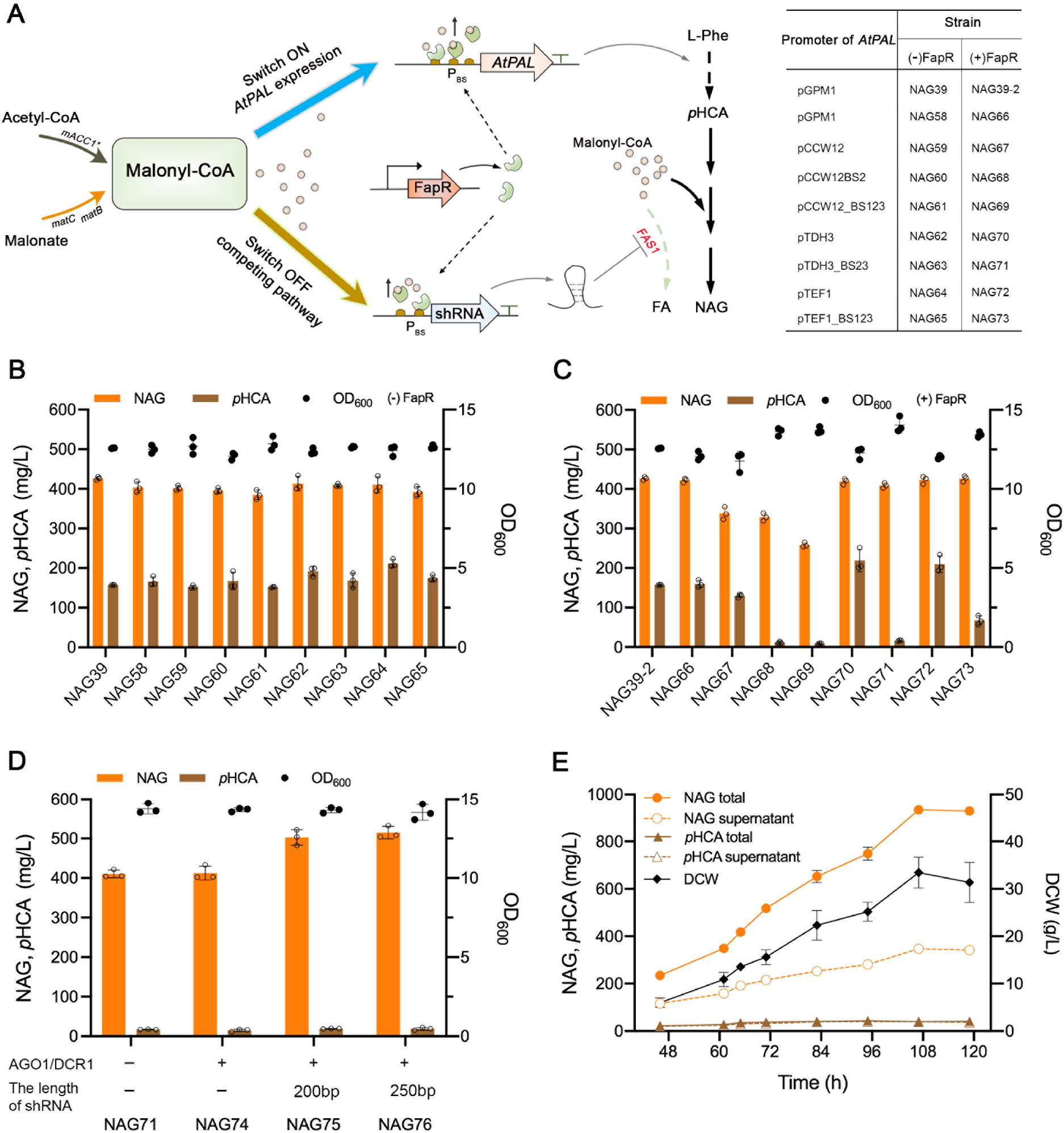
Construction of a dual dynamic control network to improve (2*S*)-naringenin production. (A) Schematic diagram of the dual dynamic control system for dynamic activation (Switch ON) and repression (Switch OFF) of genes expression. *matC*, malonate carrier protein and *matB*, malonate synthetase from *Rhizobium trifolii*; *mACC1\** represents *ACC1*^*S659A, S1157A*^, acetyl-CoA carboxylase devoid of Snf1-phosphorylation sites; *FAS1*, beta subunit of fatty acid synthetase; FA, fatty acid; L-Phe, L-phenylalanine; *p*HCA, *p*-coumaric acid; NAG, (2*S*)-naringenin. (B) Evaluation the effect of different modified promoters to control *p*-coumaric acid production on (2*S*)-naringenin production and *p*-coumaric acid accumulation in the absence of the FapR repressor. The strains with respective native promoters were used as the control. (C) Investigation of the malonyl-CoA sensor for (2*S*)-naringenin pathway regulation. Production of (2*S*)-naringenin and accumulation of *p*-coumaric acid in the strains expressed *AtPAL* under modified promoters and native promoters, were compared in presence of FapR repressor. (D) Dynamic downregulation of *FAS1* expression by RNAi under the control of the modified TDH3 promoter. (E) Fed-batch fermentation profiles of NAG76 strain in bioreactor. The biomass, total production of (2*S*)-naringenin and *p*-coumaric acid accumulation, the supernatant concentration of (2*S*)- naringenin and *p*-coumaric acid were monitored during the fed-batch cultivation. For the shake-flask fed-batch conditions, cells were grown in defined minimal medium with six tablets of FeedBeads as the sole carbon source and with 2 g/L sodium malonate dibasic supplementation. Cultures were sampled after 96 h of cultivation for metabolite analysis. All data represent the mean of n= 3 biologically independent samples and error bars show standard deviation.

To examine the activity of these modified promoters and the impact of dynamic regulation on (2*S*)- naringenin production, we constructed a series of strains by replacing the promoter of *AtPAL* in NAG39 with the modified promoters (with fap*O*, a 34 bp sequence (BS)) and the corresponding native promoters (without BS). Among the tested promoters, pGPM1, pCCW12, pTDH3 and pTEF1, led to similar production profiles and final OD even with BS inserted in the promoter region as long as FapR was not present (Fig. 4B). However, when FapR was introduced in those strains, as shown in Fig. 4C, the modified promoters exhibited different impacts on regulation of *p*-coumaric acid formation. With modified *CCW12* promoters containing one and three BS elements (in strain NAG68 and NAG69) it showed very little *p*-coumaric acid accumulation. However, the (2*S*)-naringenin production of NAG68 and NAG69 was much lower than that of NAG60 and NAG61 strains without FapR expression. Surprisingly, the NAG67 strain containing the native *CCW12* promoter displayed lower (2*S*)-naringenin production than that of NAG59 strain in absence of the FapR repressor, indicating that the basal activity of native and modified *CCW12* promoters are affected by the expression of FapR. Nevertheless, this was not the case for *TDH3* and *TEF1* promoters when compared with and without FapR expression. For the modified *TDH3* promoter, the *p*-coumaric acid accumulation in NAG71 strain was dramatically decreased to 16 mg/L, a 13-fold reduction compared with NAG70 (219 mg/L) expressing *AtPAL* under the native *TDH3* promoter. However, with the modified *TEF1* promoter in NAG73 the accumulation of *p*-coumaric acid only decreased 3-fold (67 mg/L), relative to the NAG72 control strain (209 mg/L), which may be due to the leaky expression of the modified *TEF1* promoter. Overall, these results clearly demonstrated that the dynamic regulation strategy was able to effectively reduce the accumulation of *p*-coumaric acid, whereas no increase of (2*S*)-naringenin production was found. Interestingly, when FapR was present, a significant improvement in the final biomass was observed in all the strains with modified promoters (in NAG68, NAG69, NAG71 and NAG73, Fig 4C), compared with the control strain with corresponding native promoters. This could be explained by the fact that to a certain extent the carbon flux was shifted from *p*-coumaric acid formation towards biomass production. This result inspired us to investigate whether we could further turn the saved carbon which otherwise led to *p*-coumaric acid accumulation, to the biosynthesis of (2*S*)-naringenin by further balancing *p*-coumaric acid production and malonyl-CoA availability.

### Dynamic regulation of malonyl-CoA diversion increases (2*S*)-naringenin production

In *S. cerevisiae*, the intracellular malonyl-CoA is tightly regulated and mainly consumed for synthesizing fatty acids, which are directly associated with cell growth and cell membrane biosynthesis (29). To further improve malonyl-CoA availability, we sought to fine tune the expression of fatty acid synthase through the dynamic control system combined with an RNA interference (RNAi) strategy (Fig. 4A). The fatty acid synthase complex comprises Fas1 and Fas2 subunits, and the *FAS2* gene is autoregulated by the *FAS1* gene product to coordinate the activity of the fatty acid synthase system (30). Therefore, the expression of *FAS1* was put under control of the dynamic RNAi system to divert malonyl-CoA flux from fatty acid synthesis to (2S)-naringenin biosynthesis.

To reconstitute the RNAi mechanism in *S. cerevisiae*, two heterologous genes, *AGO1* and *DCR1* (coding for Argonaute and Dicer, respectively) from *Saccharomyces castellii* (31), were firstly introduced into the genome of NAG71 strain. Small hairpin RNAs (shRNAs), consisting of both antisense and sense DNA strand of the target gene separated by a hairpin loop, have been reported to be the stronger silencing constructs compared with the double-stranded RNA generated by expression of antisense-RNA (32). Also, the hairpin length was proved to be a factor that affects the efficiency of RNAi (33). Hence, two shRNAs of different length, 200 bp and 250 bp, were designed to evaluate the efficiency of the downregulation of *FAS1*. The modified *TDH3* promoter described above, as the most tightly regulated promoter, was selected for the expression of shRNA constructs. Expressing shRNAs under the modified *TDH3* promoter further enhanced the (2*S*)-naringenin production to 502 mg/L and 514 mg/L in NAG75 and NAG76 strains, respectively, a 22% and 25% improvement compared with the control strain NAG74 (Fig. 4D). Additionally, NAG75 and NAG76 showed a final biomass similar to the NAG74 and NAG71 strains. When the shRNAs were expressed under the native *TDH3* promoter as control, all the transformants showed the formation of tiny colonies and grew very slowly, suggesting constitutive expression of shRNAs to strong silencing of fatty acid synthesis may be lethal. Overall, as a proof-of-concept, the dynamic RNAi system was proved to be functional and was able to improve (2*S*)-naringenin production.

### Fed-batch fermentation enhances (2*S*)-naringenin production

To evaluate the scalability of the dual dynamic control system, the NAG76 strain was evaluated in a bioreactor with minimal medium under glucose-limited fed-batch conditions. When glucose and ethanol were depleted at 46 h, the glucose feeding was initiated (Fig. 4E). The titer of (2*S*)-naringenin and biomass rapidly increased in an almost coupled way during the feeding process. In contrast to a rapid increase in total (2*S*)-naringenin production, the secretion of (2*S*)-naringenin was much slower, only 38% (2*S*)-naringenin was extracellular at the end of fed-batch fermentation (Fig. 4E). Finally, the highest titer of (2*S*)-naringenin reached 933 mg/L at 108 h, while the titer of *p*-coumaric acid was constantly maintained at a very low level, indicating that the dual dynamic control system could well balance the supply of two precursors and reduce the secretion of *p*-coumaric acid. However, a high amount of intracellular (2*S*)-naringenin might cause cellular toxicity on cell metabolism, requiring further strategies to increase microbial synthesis of (2*S*)-naringenin.

## Discussion

(2*S*)-Naringenin is a starting compound for synthesis of a variety of phenylpropanoid and flavonoid chemicals but also possesses many high-value pharmaceutical and nutraceutical properties. In the past decades, significant achievements have been made in microbial synthesis of (2*S*)-naringenin. Different strategies have been implemented to improve (2*S*)-naringenin production, including dynamic regulation system to balance malonyl-CoA node (3, 34), quorum-sensing based approaches for metabolic flux control (35, 36), balancing (2*S*)-naringenin biosynthesis pathway via a naringenin riboswitch (37), a promoter library (16, 38) or modular pathway engineering (15). Fewer studies, however, have focused on the spatiotemporal distribution of (2*S*)-naringenin and its intermediate *p*-coumaric acid. Here we monitored the intracellular and extracellular content of (2*S*)-naringenin and *p*-coumaric acid. We thus established a dual dynamic control system through autonomously controlling the production of *p*-coumaric acid to efficiently reduce its accumulation and downregulating a native pathway competing for malonyl-CoA to simultaneously improve (2*S*)-naringenin production.

We found that *p*-coumaric acid was easily secreted out of the yeast cell across all growth phases, regardless of high or low flux in its biosynthesis (Fig. 1C and 1D). The accumulation of *p*-coumaric acid has been observed before in *S. cerevisiae* (12) and *E. coli* (3, 15). The hypothesis was that reactions downstream of *p*-coumaric acid and/or malonyl-CoA supply was limiting (2*S*)-naringenin production. Indeed, alleviating the bottleneck downstream of *p*-coumaric acid and increasing malonyl-CoA supply both could further enhance (2*S*)-naringenin production, resulting in less *p*-coumaric acid accumulation. However, significant amounts of *p*-coumaric acid were left even at the end of fermentation (Fig.2 and 3). One may argue that if the flux towards *p*-coumaric acid was kept low there would not have this *p*-coumaric acid secretion/accumulation problem. However, to ultimately increase production of (2*S*)- naringenin and derived products, it is necessary to have high flux to its precursors. As demonstrated in Fig. 1C, increasing *p*-coumaric acid formation was able to improve (2*S*)-naringenin production.

Alternatively, the transiently accumulated *p*-coumaric acid could be potentially re-consumed later in the fermentation. This had been reported to be case, but more than one third of formed *p*-coumaric acid was not consumed at the end of fermentation (12), representing a waste of carbon. In addition, accumulation of *p*-coumaric acid was shown to be toxic for *S. cerevisiae* (39), which further points to a necessity of minimizing *p*-coumaric acid accumulation. On the other hand, re-consumption of formed *p*-coumaric acid may be hampered by inefficient import of this compound. Considering the earlier reports, supplementation of *p*-coumaric acid as substrate, the conversion yield to (2*S*)-naringenin production only reached in a range of 5.7% to 27.41% in *S. cerevisiae* even with a low *p*-coumaric acid concentration in the culture (16, 40, 41). To verify this hypothesis, we tested two background strains with different capacity in downstream of *p*-coumaric acid, IMX581N4 with only the core (2*S*)-naringenin biosynthesis pathway, and NAG41 with the enhanced downstream *p*-coumaric acid pathway and elevated malonyl-CoA supply, by exogenously feeding different concentrations of *p*-coumaric acid (100, 200, 300, 400 mg/L) (Fig. S2). With more *p*-coumaric acid supplemented, there was a general trend of increase in the total consumption. However, *p*-coumaric acid started to be accumulated when more than 200 mg/L was fed for both IMX581N4 and NAG41. This is plausible especially for NAG41, as it had a higher capacity to convert more *p*-coumaric acid to (2*S*)-naringenin. Nevertheless, the results imply the potential issue with import of *p*-coumaric acid into yeast. Surprisingly, the highest (2*S*)-naringenin production reached with addition of *p*-coumaric acid in NAG41 strain was around 76 mg/L, about one third of the corresponding NAG39 strain (214 mg/L), where the differences were with exogenous or endogenous provision of *p*-coumaric acid. These results highlighted the importance of controlling spatial content of *p*-coumaric acid.

The beneficial effect of the fed-batch model inspired us to consider dynamic control of metabolic flux as a strategy for reducing the accumulation of *p*-coumaric acid and improving (2*S*)-naringenin production. Considering that malonyl-CoA is more severely limited, we implemented a malonyl-CoA biosensor to couple *p*-coumaric acid production in response to cellular malonyl-CoA level. Fine-tuning this system resulted in almost removal of *p*-coumaric acid accumulation (Fig. 4C). Although (2*S*)-naringenin production was not improved, the cell growth (OD_600_) increased by 14% in NAG71 comparing with NAG39 using static control strategies. To channel carbon flux toward (2*S*)-naringenin synthesis we further combined this malonyl-CoA sensor with an RNAi strategy to control fatty acid biosynthesis. This dual dynamic system further increased (2*S*)-naringenin by 25%, without significant accumulation of *p*-coumaric acid and any effect on cell growth (Fig. 4D). Recently, a growth-coupled NCOMB (Naringenin-Coumaric acid-Malonyl-CoA-Balanced) dynamic regulation network was reported to successfully improve (2*S*)-naringenin production with a concomitant increase in cell growth and reduced accumulation of *p*-coumaric acid compared to the strain using static strain engineering approach in *E. coli* (3). However, this system requires the presence of intracellular (2*S*)-naringenin and *p*-coumaric acid, which limits its application in other organisms like *S. cerevisiae* and prevents complete removal of *p*-coumaric acid accumulation.

Finally, a (2*S*)-naringenin titer of 933 mg/L was achieved from about 128 g/L of glucose in minimal medium in about 108 h (Fig. 4E). Considering the difference in cultivation conditions and strategies, this production level is comparable with *de novo* (2*S*)-naringenin biosynthesis in other microorganisms, i.e., 898 mg/L (2*S*)-naringenin was generated in *Y. lipolytica* from minimal medium with 180 g/L glucose in about 300 h (8); 1129.44 mg/L (2*S*)-naringenin was achieved in yeast in about 108 h by using rich medium YPD with more than 187 g/L glucose (11). However, our study showed higher carbon yield and productivity and provided an effective strategy to address *p*-coumaric acid accumulation. We believe that the methods developed here could be relevant for other microbial cell factories for producing plant natural products such as flavonoids and alkaloids. Furthermore, the spatiotemporal control of intermediates could be extremely important for developing co-culture systems for microbial production.

## Materials and Methods

### Strains and reagents

All the *S. cerevisiae* strains and plasmids used in this work are listed in Supplementary Table 1 and 2. High-fidelity Phusion DNA polymerase was purchased from New England Biolabs. PrimeStar DNA polymerase and SapphireAmp^*^ Fast PCR Master Mix were purchased from TaKaRa Bio. Plasmid extraction and DNA gel purification kits were purchased from ThermoFisher Scientific. All oligonucleotides were synthesized at IDT Biotechnology company and are listed in Supplementary Table 3. All codon-optimized heterologous genes were synthesized at Genscript and are listed in Supplementary Table 4. All chemicals including analytical standards were purchased from Sigma-Aldrich.

### Strain cultivation

*S. cerevisiae* strains for preparation of competent cells were cultivated in YPD medium consisting of 10 g/L yeast extract (Merck Millipore), 20 g/L peptone (Difco) and 20 g/L glucose (Merck Millipore). Synthetic complete medium without uracil (SC-URA), which consisted of 6.7 g/L yeast nitrogen base without amino acids (Formedium), 0.77 g/L CSM without uracil (Formedium), 20 g/L glucose (Merck Millipore) and 20 g/L agar (Merck Millipore), was used for selection of yeast transformants containing *URA3* marker-based plasmids. To remove the *URA3* maker, yeast transformants were selected against on synthetic complete medium with 5-fluoroorotic acid plates (SC+5-FOA), which contained 6.7 g/L yeast nitrogen base, 0.77 g/L CSM and 0.8 g/L 5-FOA.

Shake flask batch fermentations for the production of (2*S*)-naringenin and *p*-coumaric acid were carried out in minimal medium, containing 7.5 g/L (NH_4_)_2_SO_4_, 14.4 g/L KH_2_PO_4_, 0.5 g/L MgSO_4_·7H_2_O, and 30 g/L glucose, 2 mL/L trace metal (3.0 g/L FeSO_4_·7H_2_O, 4.5 g/L ZnSO_4_·7H_2_O, 4.5 g/L CaCl_2_·2H_2_O, 0.84 g/L MnCl_2_·2H_2_O, 0.3 g/L CoCl_2_·6H_2_O, 0.3 g/L CuSO_4_·5H_2_O, 0.4 g/L Na_2_MoO_4_·2H_2_O, 1.0 g/L H_3_BO_3_, 0.1 g/L KI, and 19.0 g/L Na_2_EDTA·2H_2_O), and 1mL/L vitamin solutions (0.05 g/L D-biotin, 1.0 g/L D-pantothenic acid hemicalcium salt, 1.0 g/L thiamin–HCl, 1.0 g/L pyridoxin–HCl, 1.0 g/L nicotinic acid, 0.2 g/L 4-aminobenzoic acid, and 25.0 g/L myo-inositol) supplemented with 60 mg/L uracil if needed. Three biological replicates of each engineered strain were inoculated in tubes with 2 mL minimal medium at 30 °C with 220 rpm agitation for 24 h. Then, the precultures were inoculated to the initial OD600 of 0.05 in 15 mL minimal medium or minimal medium containing 2 g/L sodium malonate dibasic (Sigma-Aldrich) in 100 mL unbaffled shake-flasks. The cells were cultured at 30 °C with 220 rpm agitation for 96 h. For shake flask fermentation in mimicked fed-batch condition, six tablets of FeedBeads (SMFB08001, Kuhner Shaker, Basel, Switzerland), corresponding to 30 g/L glucose, were used as the sole carbon source and cultivations were run for 96 h at 30 °C with 220 rpm agitation.

The fed-batch bioreactor fermentation was performed in biological quadruplicates in 1 L bioreactors (DasGip Parallel Bioreactor System, DasGip, Germany) containing 250 mL initial minimal medium with 30 g/L glucose. The temperature (30 °C), pH, agitation and aeration were monitored and controlled using a DasGip Control 4.0 System. The dissolved oxygen level was maintained above 30%. The pH was set at 5.6 and controlled by the addition of 4 M KOH and 2 M HCl. The agitation and aeration were initially set to 600 rpm and 36 L/h respectively. When the dissolved oxygen in the reactor decreased below 30%, the agitation increased to 1000 rpms and the airflow to 48 L/h. The aeration was controlled and provided by a DasGip MX4/4 module. The composition of the off-gas was monitored using a DasGip Off Gas Analyzer GA4. Addition of the acid, base, and glucose feeding was conducted with DasGip MP8 multi-pump modules (pump head tubing: 0.5 mm ID, 1.0 mm wall thickness). During the fed-batch cultivation, the cells were fed a 200 g/L glucose solution with a feeding rate that was exponentially increased (μ = 0.05 h^−1^) to maintain a constant growth rate. The used minimal medium contained: 15 g/L (NH_4_)_2_SO_4_, 9 g/L KH_2_PO_4_, 1.5 g/L MgSO_4_·7H_2_O, 180 mg/L uracil, 3 ml/L trace metal, 3 ml/L vitamin solutions and 9 g/L sodium malonate dibasic. The initial feeding rate was calculated by using the biomass yield and concentration that were obtained during prior duplicate batch cultivations with this strain. The feeding was initiated after the CO_2_ levels dropped indicating that glucose and ethanol were totally consumed. Cell dry weight (CDW) measurements were performed by filtrating 1 mL of cell culture through a weighed 0.45 μm filter membrane (Sartorius Biolab, Gottingen, Germany) and weighting the filter after microwaving it for 15 min and letting it dry in a desiccator for 24h.

### DNA manipulation

The *S. cerevisiae* CEN.PK113-5D-detivative IMX581 (*MATa ura3-52 can1Δ::cas9-natNT2 TRP1 LEU2 HIS3*)(42) was used as original strain for all engineering work. CRISPR–Cas9-mediated genome engineering was used for chromosome-based gene overexpression and the integration of expression cassettes. IMX581 genomic DNA served as the template for all native promoters, genes and terminators. For optimized heterologous genes, synthetic fragments or plasmids were utilized as the DNA template for amplification. High-fidelity Phusion DNA polymerase was used for entire DNA fragment amplifications, except PrimeSTAR HS polymerase was selected for *in vitro* fusion PCR for generating integration modules. Functional expression modules were generated according to the overlapping extension PCR procedure (43), and all used integration modules are listed in Supplementary Table 5.

Specific chromosomal loci (44), enabling stable and high-level expression of heterologous genes, were selected as the recombination sites for integration of gene overexpression modules. Taking construction of NAG05 strain as an example to show the CRISPR/Cas9-mediated integration of in-vitro-assembled gene expression modules, the integration module was divided into two fragments: M1 (*XII-4 up-TDH3p-At4CL1-ADH1t +TDH2t*) and M6 (*ADH1t+TDH2t-RsCHS-CCW12p+tHXT7p-PsCHI-FBAt-XII-4 dn*). M1 was generated by fusing DNA parts including *XII-4 up, TDH3p, At4CL, ADH1t, TDH2t*. Specifically, the upstream homologous arm *XII-4 up* (P001, P002), promoters *TDH3p* (P045, P046), terminators *ADH1t* (P047, P048) and *TDH2t* (P049, P050) were amplified from IMX581 genomic DNA, respectively; codon-optimized gene *At4CL1* (P164, P165) was amplified from pCfB0854. M6 was assembled by fusing the DNA parts of *ADH1t, TDH2t, RsCHS, CCW12p, tHXT7p, PsCHI, FBAt* and *XII-4 dn*. Codon-optimized genes *RsCHS* (P166, P167) and *PsCHI* (P168, P169) were amplified by using the synthetic genes as template. Promoters *CCW12p* (P051, P052) and *tHXT7p* (P053, P054), terminator *FBAt* (P055, P056) and the downstream homologous arm *XII-4 dn* (P003, P004) were amplified from IMX581 genomic DNA, respectively. Then, the equimolar amounts of purified fragments M1 and M6 (50-100 ng/kb) with gRNA plasmid pQC010 were co-transformed into the IMX581 strain by using the lithium acetate-mediated yeast transformation protocol (42) and transformants selected on SC-URA plates. The grown clones were verified by colony PCR by using SapphireAmp^®^ Fast PCR Master Mix. Subsequently, the correct transformants were streaked on SC+5-FOA plates to loop out the gRNA plasmid and recycle the *URA3* marker.

The *FAS1* hairpin part was constructed according to a described method (32), the expression module was divided into two DNA fragments: the sense region was assembled by fusing the DNA parts of upstream homologous fragments of the chromosomal target site, promoter, 250bp sense sequence of the target gene and the 80-bp sequence spanning intron 1 from *Schizosaccharomyces pombe rad9* representing a hairpin loop. The antisense region was generated through assembling the DNA parts of 80bp sequence of intron 1 from *S. pombe rad9*, the antisense region of the target gene (reverse complement of sense region), terminator and downstream homologous fragments of the chromosomal target site. Subsequently, the two fragments were co-transformed and assembled into the genomic integration site via *in vivo* homologous recombination. The correct transformants were then confirmed by colony PCR amplification.

To construct specific guide RNAs for a selected gene/genomic locus, all potential gRNAs were selected and compared with all potential off-targets in the whole CEN.PK113-7D genome using the CRISPRdirect webtool (http://crispr.dbcls.jp/) (45). The gRNA plasmids were constructed according to the Gibson assembly method in which gRNA sequence-containing DNA parts were *in vitro* recombined with a vector backbone (42). The correct recombinant plasmids were verified by *E. coli* colony PCR and further confirmed by DNA sequencing.

### Metabolite extraction and quantification

For total production analysis of (2*S*)-naringenin and *p*-coumaric acid, 0.5 mL of the culture samples were thoroughly mixed with an equal volume of absolute ethanol (100% v/v) and centrifuged at 12,000 rpm for 10 min. The supernatants were then collected and stored in -20 °C until HPLC detection. To measure the supernatant concentrations of (2*S*)-naringenin and *p*-coumaric acid, a 1.0 mL culture sample was centrifuged at 12,000 rpm for 10 min and filtered through 0.45 μm syringe filter. The filtered supernatants were then saved for HPLC analysis.

Samples were quantified on a Dionex Ultimate 3000 HPLC (Thermo Fisher Scientific, Waltham, MA, USA) equipped with a Discovery HS F5 150 mm × 46 mm column (particle size 5 μm) (Sigma-Aldrich, St. Louis, MO, USA). The column was kept at 30 °C, and metabolites from 10 μL of supernatants were separated. The HPLC analysis was carried out with 10 mM ammonium formate (pH 3.0, adjusted by formic acid) (Solvent A) and acetonitrile (Solvent B) as the eluents. The eluent flow rate was 1.5 mL/min. The gradient program started with 5% of solvent B (0–0.5 min), increased linearly from 5% to 60% (0.5– 9.5 min), then the second linear increase from 60% to 100% (9.5–10.5 min) and maintained at 100% for 0.5 min (10.5–11 min), at last a linear decrease from 100% to 5% (11–11.5 min) and maintained at 5% for 1.5 min (11.5–13 min). *p*-Coumaric acid was detected by absorbance at 304 nm with a retention time of 6.3 min. (2*S*)-naringenin was detected by absorbance at 289 nm with a retention time of 8.9 min. The (2*S*)-naringenin and *p*-coumaric acid concentrations were calculated from standard curves.

For residual glucose detection, the collected culture samples were centrifuged at 12,000 rpm for 10 min and filtered through 0.45 μm syringe filter. The supernatants were then quantified by HPLC (Thermo Fisher Scientific, CA, USA) equipped with an Aminex HPX-87G column (Bio-Rad, Hercules, CA, USA) and a UV and RI detector. 5 mM H_2_SO_4_ was used as the mobile phase and the column was kept at 45 °C with a flow rate of 0.6 mL/min for 35 min. The glucose concentrations were calculated from the standard curves.

## Acknowledgments

We acknowledge funding from the European Union’s Horizon 2020 research and innovation program under Grant Agreement No. 686070, the Novo Nordisk Foundation (grant no. NNF10CC1016517 and NNF18OC0034844). We also acknowledge funding support from Vetenskapsrådet and Stiftelsen för internationalisering av högre utbildning och forskning.

## Contributions

Y.C. and J.M. designed the study; J.M., M.T.M., X.L., and Q.L. performed the experiments; J.M., M.T.M., X.L., Q.L., J.N., V.S., and Y.C. analyzed the data; J.M. and Y.C. wrote the paper and all authors helped revising and approved the final version.

