## Supplementary for "Fine-tuning of coumaric acid synthesis to increase naringenin production in yeast"

**Figure S1.**

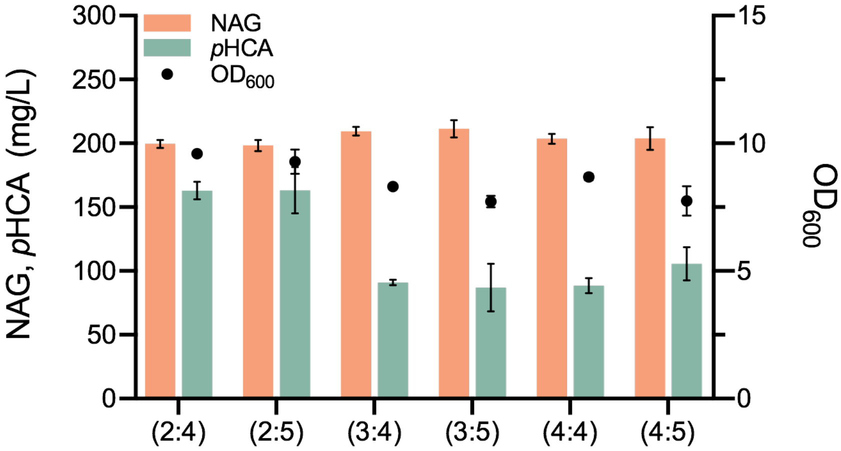


**Figure S1. Optimization of the downstream enzymes of *p*-coumaric acid for (2*S*)-naringenin production.** The numbers (n : n) indicate the respective gene copy numbers of *4CL* and *CHS*&*CHI*. Cells were grown in a defined minimal medium with 30 g/L glucose as the sole carbon source, and cultures were sampled after 96 h of growth for metabolite analysis. All data represent the mean of n = 3 biologically independent samples and error bars show standard deviation.

**Figure S2**

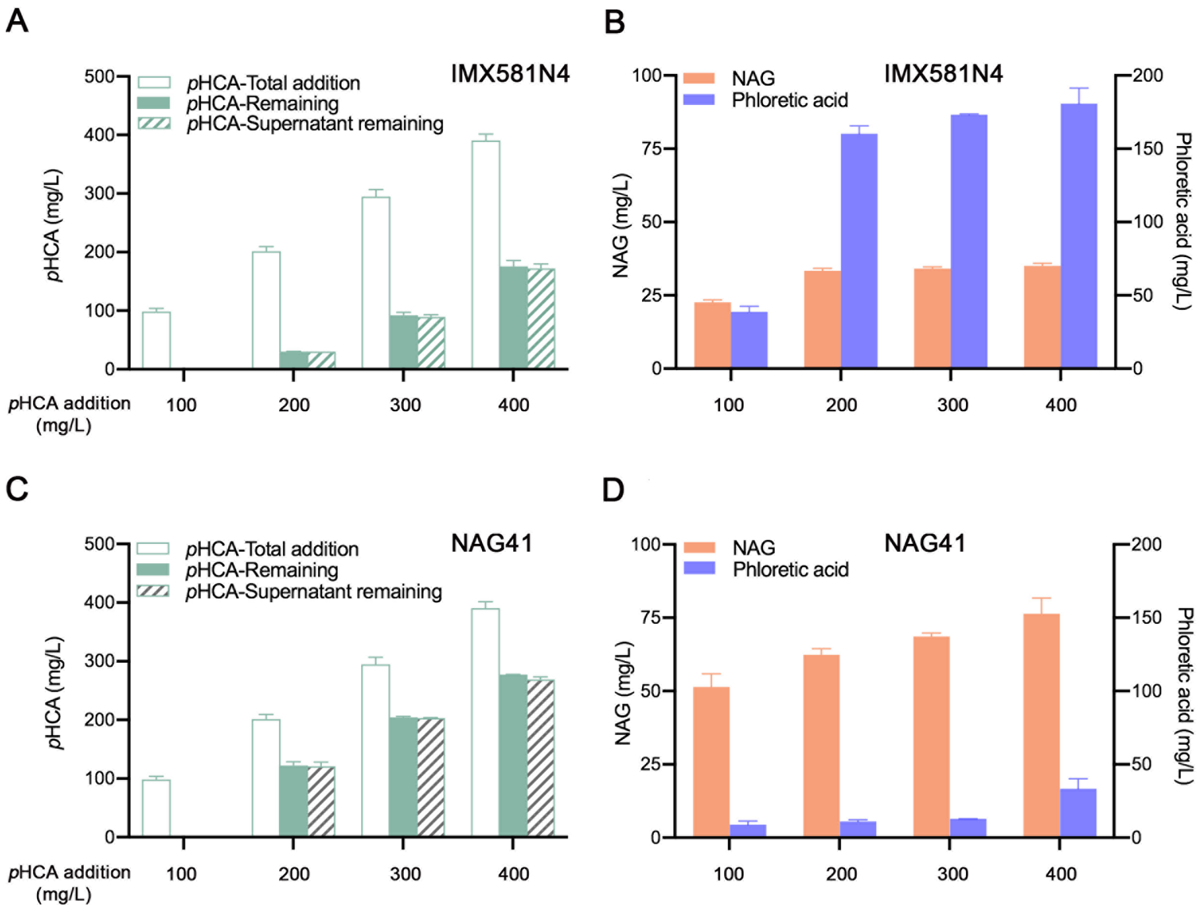


**Figure S2. Production of (2*S*)-naringenin in strains supplemented with different concentrations of *p*-coumaric acid.** (A) (B) (2*S*)-naringenin production by the IMX581N4 strain expressing one copy of *4CL* and *CHS*&*CHI* with the supplementary of different concentrations of *p*-coumaric acid. Cells were grown in a defined minimal medium with 30 g/L glucose as the sole carbon source and with supplemented *p*-coumaric acid (100, 200, 300, 400 mg/L) as precursor. Cultures were sampled after 96 h of growth for metabolite detection. (C) (D) Production of (2*S*)-naringenin in strain NAG41 supplemented with different concentrations of *p*-coumaric acid. Strain NAG41 harboring three copies of *4CL* and four copies of *CHS*&*CHI*, simultaneously expressing the *ACC1* mutant and the malonate assimilation pathway (*matBC*). Cells were grown in a defined minimal medium with 30 g/L glucose as the sole carbon source and 2 g/L sodium malonate dibasic, and with supplemented *p*-coumaric acid (100, 200, 300, 400 mg/L) as precursor. Cultures were sampled after 96 h of growth for metabolite analysis. All data represent the mean of n ≥ 2 biologically independent samples and error bars show standard deviation.

### Supplementary Table 1. *S. cerevisiae* strains used in this study.

| Strain ID | Genotype | Parental strain | Origin |
|---|---|---|---|
| IMX581 | *MATa ura3-52 can1Δ :: cas9-natNT2 TRP1 LEU2 HIS3* | | (1) |
| QL01 | *MATa ura3-52 can1Δ::cas9-natNT2 TRP1 LEU2 HIS3 XII-2::(GPM1p-AtPAL2-FBA1t)+(TDH3p-AtC4H-CYC1t)+(tHXT7p-AtATR2-pYX212t)+(PGK1p-CYB5-ADH1t)* | IMX581 | (2) |
| NAG01 | *MATa ura3-52 can1Δ::cas9-natNT2 TRP1 LEU2 HIS3 XII-2::(GPM1p-AtPAL2-FBA1t)+(TDH3p-AtC4H-CYC1t)+(tHXT7p-AtATR2-pYX212t)+(PGK1p-CYB5-ADH1t) XII-4::(TDH3p-At4CL-ADH1t) + (TDH2t-HaCHS-CCW12p)+(tHXT7p-PhCHI-FBAt)* | QL01 | This work |
| NAG02 | *MATa ura3-52 can1Δ::cas9-natNT2 TRP1 LEU2 HIS3 XII-2::(GPM1p-AtPAL2-FBA1t)+(TDH3p-AtC4H-CYC1t)+(tHXT7p-AtATR2-pYX212t)+(PGK1p-CYB5-ADH1t) XII-4::(TDH3p-At4CL-ADH1t) + (TDH2t-HaCHS-CCW12p)+(tHXT7p-PsCHI-FBAt)* | QL01 | This work |
| NAG03 | *MATa ura3-52 can1Δ::cas9-natNT2 TRP1 LEU2 HIS3 XII-2::(GPM1p-AtPAL2-FBA1t)+(TDH3p-AtC4H-CYC1t)+(tHXT7p-AtATR2-pYX212t)+(PGK1p-CYB5-ADH1t) XII-4::(TDH3p-At4CL-ADH1t) + (TDH2t-HaCHS-CCW12p)+(tHXT7p-SmCHI-FBAt)* | QL01 | This work |
| NAG04 | *MATa ura3-52 can1Δ::cas9-natNT2 TRP1 LEU2 HIS3 XII-2::(GPM1p-AtPAL2-FBA1t)+(TDH3p-AtC4H-CYC1t)+(tHXT7p-AtATR2-pYX212t)+(PGK1p-CYB5-ADH1t) XII-4::(TDH3p-At4CL-ADH1t) + (TDH2t-RsCHS-CCW12p)+(tHXT7p-PhCHI-FBAt)* | QL01 | This work |
| NAG05 | *MATa ura3-52 can1Δ::cas9-natNT2 TRP1 LEU2 HIS3 XII-2::(GPM1p-AtPAL2-FBA1t)+(TDH3p-AtC4H-CYC1t)+(tHXT7p-AtATR2-pYX212t)+(PGK1p-CYB5-ADH1t) XII-4::(TDH3p-At4CL-ADH1t) + (TDH2t-RsCHS-CCW12p)+(tHXT7p-PsCHI-FBAt)* | QL01 | This work |
| NAG06 | *MATa ura3-52 can1Δ::cas9-natNT2 TRP1 LEU2 HIS3 XII-2::(GPM1p-AtPAL2-FBA1t)+(TDH3p-AtC4H-CYC1t)+(tHXT7p-AtATR2-pYX212t)+(PGK1p-CYB5-ADH1t) XII-4::(TDH3p-At4CL-ADH1t) + (TDH2t-RsCHS-CCW12p)+(tHXT7p-SmCHI-FBAt)* | QL01 | This work |
| NAG07 | *MATa ura3-52 can1Δ::cas9-natNT2 TRP1 LEU2 HIS3 XII-2::(GPM1p-AtPAL2-FBA1t)+(TDH3p-AtC4H-CYC1t)+(tHXT7p-AtATR2-pYX212t)+(PGK1p-CYB5-ADH1t) XII-4:: (TDH3p-At4CL-ADH1t) + (TDH2t-SmCHS-CCW12p)+(tHXT7p-PhCHI-FBAt)* | QL01 | This work |
| NAG08 | *MATa ura3-52 can1Δ::cas9-natNT2 TRP1 LEU2 HIS3 XII-2::(GPM1p-AtPAL2-FBA1t)+(TDH3p-AtC4H-CYC1t)+(tHXT7p-AtATR2-pYX212t)+(PGK1p-CYB5-ADH1t) XII-4::(TDH3p-At4CL-ADH1t) + (TDH2t-SmCHS-CCW12p)+(tHXT7p-PsCHI-FBAt)* | QL01 | This work |
| NAG09 | *MATa ura3-52 can1Δ::cas9-natNT2 TRP1 LEU2 HIS3 XII-2::(GPM1p-AtPAL2-FBA1t)+(TDH3p-AtC4H-CYC1t)+(tHXT7p-AtATR2-pYX212t)+(PGK1p-CYB5-ADH1t) XII-4::(TDH3p-At4CL-ADH1t) + (TDH2t-SmCHS-CCW12p)+(tHXT7p-SmCHI-FBAt)* | QL01 | This work |
| NAG10 | *MATa ura3-52 can1Δ::cas9-natNT2 TRP1 LEU2 HIS3 XII-2::(GPM1p-AtPAL2-FBA1t)+(TDH3p-AtC4H-CYC1t)+(tHXT7p-AtATR2-pYX212t)+(PGK1p-CYB5-ADH1t) X-3::(TPI1p-EcaroL-pYX212t)+(ADH1t-ARO7* $^{G141S}$*-TEF1p)+(PGK1p-ARO4* $^{K229L}$*-CYC1t) X-4::(CYC1t-ARO1-TPI1p)+(TDH3p-ARO2-ADH1t)+(TDH2t-ARO3-TEF1p) XII-4::(TDH3p-At4CL-ADH1t) + (TDH2t-RsCHS-CCW12p)+(tHXT7p-PsCHI-FBAt)* | NAG05 | This work |
| NAG11 | *MATa ura3-52 can1Δ::cas9-natNT2 TRP1 LEU2 HIS3 XII-2::(GPM1p-AtPAL2-FBA1t)+(TDH3p-AtC4H-CYC1t)+(tHXT7p-AtATR2-pYX212t)+(PGK1p-CYB5-ADH1t) X-3::(TPI1p-EcaroL-pYX212t)+(ADH1t-ARO7* $^{G141S}$*-TEF1p)+(PGK1p-ARO4* $^{K229L}$*-CYC1t) X-4::(CYC1t-ARO1-TPI1p)+(TDH3p-ARO2-ADH1t)+(TDH2t-ARO3-TEF1p) X-2::( GPM1p-PHA2-CYC1t) XII-4::(TDH3p-At4CL-ADH1t) + (TDH2t-RsCHS-CCW12p)+(tHXT7p-PsCHI-FBAt)* | NAG010 | This work |
| NAG1-2 | *MATa ura3-52 can1Δ::cas9-natNT2 TRP1 LEU2 HIS3 XII-2::(GPM1p-AtPAL2-FBA1t)+(TDH3p-AtC4H-CYC1t)+(tHXT7p-AtATR2-pYX212t)+(PGK1p-CYB5-ADH1t) X-3::(TPI1p-EcaroL-pYX212t)+(ADH1t-ARO7* $^{G141S}$*-TEF1p)+(PGK1p-ARO4* $^{K229L}$*-CYC1t) X-4::(CYC1t-ARO1-TPI1p)+(TDH3p-ARO2-ADH1t)+(TDH2t-ARO3-TEF1p) XII-4::(TDH3p-At4CL-ADH1t) + (TDH2t-RsCHS-* | NAG010 | This work |

| | | | |
|---|---|---|---|
| | *CCW12p)+(tHXT7p-PsCHI-FBAt) XII-5::(pYX212t-PsCHI-PGKp)+ (TEF1p-RsCHS-FBAt)* | | |
| NAG1-3 | *MATa ura3-52 can1Δ::cas9-natNT2 TRP1 LEU2 HIS3 XII-2::(GPM1p-AtPAL2-FBA1t)+(TDH3p-AtC4H-CYC1t)+(tHXT7p-AtATR2-pYX212t)+(PGK1p-CYB5-ADH1t) X-3::(TPI1p-EcaroL-pYX212t)+(ADH1t-ARO7* $^{G141S}$*-TEF1p)+(PGK1p-ARO4* $^{K229L}$*-CYC1t) X-4::(CYC1t-ARO1-TPI1p)+(TDH3p-ARO2-ADH1t)+(TDH2t-ARO3-TEF1p) XII-4::(TDH3p-At4CL-ADH1t) + (TDH2t-RsCHS-CCW12p)+(tHXT7p-PsCHI-FBAt) XII-5::(pYX212t-PsCHI-PGKp)+ (TEF1p-RsCHS-FBAt) XI-1::(pYX212t-PsCHI-PGKp)+ (TEF1p-RsCHS-FBAt)* | NAG1-2 | This work |
| NAG1-4 | *MATa ura3-52 can1Δ::cas9-natNT2 TRP1 LEU2 HIS3 XII-2::(GPM1p-AtPAL2-FBA1t)+(TDH3p-AtC4H-CYC1t)+(tHXT7p-AtATR2-pYX212t)+(PGK1p-CYB5-ADH1t) X-3::(TPI1p-EcaroL-pYX212t)+(ADH1t-ARO7* $^{G141S}$*-TEF1p)+(PGK1p-ARO4* $^{K229L}$*-CYC1t) X-4::(CYC1t-ARO1-TPI1p)+(TDH3p-ARO2-ADH1t)+(TDH2t-ARO3-TEF1p) XII-4::(TDH3p-At4CL-ADH1t) + (TDH2t-RsCHS-CCW12p)+(tHXT7p-PsCHI-FBAt) XII-5::(pYX212t-PsCHI-PGKp)+ (TEF1p-RsCHS-FBAt) XI-1::(pYX212t-PsCHI-PGKp)+ (TEF1p-RsCHS-FBAt) XII-1::(TDH2t-RsCHS-CCW12p)+(tHXT7p-PsCHI-FBAt)* | NAG1-3 | This work |
| NAG2-1 | *MATa ura3-52 can1Δ::cas9-natNT2 TRP1 LEU2 HIS3 XII-2::(GPM1p-AtPAL2-FBA1t)+(TDH3p-AtC4H-CYC1t)+(tHXT7p-AtATR2-pYX212t)+(PGK1p-CYB5-ADH1t) X-3::(TPI1p-EcaroL-pYX212t)+(ADH1t-ARO7* $^{G141S}$*-TEF1p)+(PGK1p-ARO4* $^{K229L}$*-CYC1t) X-4::(CYC1t-ARO1-TPI1p)+(TDH3p-ARO2-ADH1t)+(TDH2t-ARO3-TEF1p) XII-4::(TDH3p-At4CL-ADH1t) + (TDH2t-RsCHS-CCW12p)+(tHXT7p-PsCHI-FBAt) XII-1::(TDH3p-At4CL-ADH1t)* | NAG010 | This work |
| NAG2-2 | *MATa ura3-52 can1Δ::cas9-natNT2 TRP1 LEU2 HIS3 XII-2::(GPM1p-AtPAL2-FBA1t)+(TDH3p-AtC4H-CYC1t)+(tHXT7p-AtATR2-pYX212t)+(PGK1p-CYB5-ADH1t) X-3::(TPI1p-EcaroL-pYX212t)+(ADH1t-ARO7* $^{G141S}$*-TEF1p)+(PGK1p-ARO4* $^{K229L}$*-CYC1t) X-4::(CYC1t-ARO1-TPI1p)+(TDH3p-ARO2-ADH1t)+(TDH2t-ARO3-TEF1p) XII-4:: (TDH3p-At4CL-ADH1t) + (TDH2t-RsCHS-CCW12p)+(tHXT7p-PsCHI-FBAt) XII-1::(TDH3p-At4CL-ADH1t) + (TDH2t-RsCHS-CCW12p)+(tHXT7p-PsCHI-FBAt)* | NAG10 | This work |
| NAG2-3 | *MATa ura3-52 can1Δ::cas9-natNT2 TRP1 LEU2 HIS3 XII-2::(GPM1p-AtPAL2-FBA1t)+(TDH3p-AtC4H-CYC1t)+(tHXT7p-AtATR2-pYX212t)+(PGK1p-CYB5-ADH1t) X-3::(TPI1p-EcaroL-pYX212t)+(ADH1t-ARO7* $^{G141S}$*-TEF1p)+(PGK1p-ARO4* $^{K229L}$*-CYC1t) X-4::(CYC1t-ARO1-TPI1p)+(TDH3p-ARO2-ADH1t)+(TDH2t-ARO3-TEF1p) XII-4::(TDH3p-At4CL-ADH1t) + (TDH2t-RsCHS-CCW12p)+(tHXT7p-PsCHI-FBAt) XII-1::(TDH3p-At4CL-ADH1t) + (TDH2t-RsCHS-CCW12p)+(tHXT7p-PsCHI-FBAt) XII-5::(pYX212t-PsCHI-PGKp)+ (TEF1p-RsCHS-FBAt)* | NAG2-2 | This work |
| NAG2-4 | *MATa ura3-52 can1Δ::cas9-natNT2 TRP1 LEU2 HIS3 XII-2::(GPM1p-AtPAL2-FBA1t)+(TDH3p-AtC4H-CYC1t)+(tHXT7p-AtATR2-pYX212t)+(PGK1p-CYB5-ADH1t) X-3::(TPI1p-EcaroL-pYX212t)+(ADH1t-ARO7* $^{G141S}$*-TEF1p)+(PGK1p-ARO4* $^{K229L}$*-CYC1t) X-4::(CYC1t-ARO1-TPI1p)+(TDH3p-ARO2-ADH1t)+(TDH2t-ARO3-TEF1p) XII-4::(TDH3p-At4CL-ADH1t) + (TDH2t-RsCHS-CCW12p)+(tHXT7p-PsCHI-FBAt) XII-1::(TDH3p-At4CL-ADH1t) + (TDH2t-RsCHS-CCW12p)+(tHXT7p-PsCHI-FBAt) XII-5::(pYX212t-PsCHI-PGKp)+ (TEF1p-RsCHS-FBAt) XI-1::(pYX212t-PsCHI-PGKp)+ (TEF1p-RsCHS-FBAt)* | NAG2-3 | This work |
| NAG3-1 | *MATa ura3-52 can1Δ::cas9-natNT2 TRP1 LEU2 HIS3 XII-2::(GPM1p-AtPAL2-FBA1t)+(TDH3p-AtC4H-CYC1t)+(tHXT7p-AtATR2-pYX212t)+(PGK1p-CYB5-ADH1t) X-3::(TPI1p-EcaroL-pYX212t)+(ADH1t-ARO7* $^{G141S}$*-TEF1p)+(PGK1p-ARO4* $^{K229L}$*-CYC1t) X-4::(CYC1t-ARO1-TPI1p)+(TDH3p-ARO2-ADH1t)+(TDH2t-ARO3-TEF1p) XII-4::(TDH3p-At4CL-ADH1t) + (TDH2t-RsCHS-CCW12p)+(tHXT7p-PsCHI-FBAt) XII-1::(TDH3p-At4CL-ADH1t) XII-5::(CYC1t-At4CL-TPIp)* | NAG2-1 | This work |
| NAG3-2 | *MATa ura3-52 can1Δ::cas9-natNT2 TRP1 LEU2 HIS3 XII-2::(GPM1p-AtPAL2-FBA1t)+(TDH3p-AtC4H-CYC1t)+(tHXT7p-AtATR2-pYX212t)+(PGK1p-CYB5-ADH1t) X-3::(TPI1p-EcaroL-pYX212t)+(ADH1t-ARO7* $^{G141S}$*-TEF1p)+(PGK1p-ARO4* $^{K229L}$*-CYC1t) X-4::(CYC1t-ARO1-TPI1p)+(TDH3p-ARO2-ADH1t)+(TDH2t-ARO3-TEF1p) XII-4::(TDH3p-At4CL-ADH1t) + (TDH2t-RsCHS-CCW12p)+(tHXT7p-PsCHI-FBAt) XII-1::(TDH3p-At4CL-ADH1t) + (TDH2t-RsCHS-CCW12p)+(tHXT7p-PsCHI-FBAt) XII-5::(CYC1t-At4CL-TPIp)* | NAG2-2 | This work |

| | | | |
|---|---|---|---|
| NAG3-3 | *MATa ura3-52 can1Δ::cas9-natNT2 TRP1 LEU2 HIS3 XII-2::(GPM1p-AtPAL2-FBA1t)+(TDH3p-AtC4H-CYC1t)+(tHXT7p-AtATR2-pYX212t)+(PGK1p-CYB5-ADH1t) X-3::(TPI1p-EcaroL-pYX212t)+(ADH1t-ARO7* $^{G141S}$*-TEF1p)+(PGK1p-ARO4* $^{K229L}$*-CYC1t) X-4::(CYC1t-ARO1-TPI1p)+(TDH3p-ARO2-ADH1t)+(TDH2t-ARO3-TEF1p) XII-4::(TDH3p-At4CL-ADH1t) + (TDH2t-RsCHS-CCW12p)+(tHXT7p-PsCHI-FBAt) XII-1::(TDH3p-At4CL-ADH1t) + (TDH2t-RsCHS-CCW12p)+(tHXT7p-PsCHI-FBAt) XII-5::(pYX212t-PsCHI-PGKp)+(TEF1p-RsCHS-FBAt) + (CYC1t-At4CL-TPIp)* | NAG2-2 | This work |
| NAG3-4 | *MATa ura3-52 can1Δ::cas9-natNT2 TRP1 LEU2 HIS3 XII-2::(GPM1p-AtPAL2-FBA1t)+(TDH3p-AtC4H-CYC1t)+(tHXT7p-AtATR2-pYX212t)+(PGK1p-CYB5-ADH1t) X-3::(TPI1p-EcaroL-pYX212t)+(ADH1t-ARO7* $^{G141S}$*-TEF1p)+(PGK1p-ARO4* $^{K229L}$*-CYC1t) X-4::(CYC1t-ARO1-TPI1p)+(TDH3p-ARO2-ADH1t)+(TDH2t-ARO3-TEF1p) XII-4::(TDH3p-At4CL-ADH1t) + (TDH2t-RsCHS-CCW12p)+(tHXT7p-PsCHI-FBAt) XII-1::(TDH3p-At4CL-ADH1t) + (TDH2t-RsCHS-CCW12p)+(tHXT7p-PsCHI-FBAt) XII-5::(pYX212t-PsCHI-PGKp)+(TEF1p-RsCHS-FBAt) + (CYC1t-At4CL-TPIp) XI-1::(pYX212t-PsCHI-PGKp)+(TEF1p-RsCHS-FBAt)* | NAG3-3 | This work |
| NAG4-1 | *MATa ura3-52 can1Δ::cas9-natNT2 TRP1 LEU2 HIS3 XII-2::(GPM1p-AtPAL2-FBA1t)+(TDH3p-AtC4H-CYC1t)+(tHXT7p-AtATR2-pYX212t)+(PGK1p-CYB5-ADH1t) X-3::(TPI1p-EcaroL-pYX212t)+(ADH1t-ARO7* $^{G141S}$*-TEF1p)+(PGK1p-ARO4* $^{K229L}$*-CYC1t) X-4::(CYC1t-ARO1-TPI1p)+(TDH3p-ARO2-ADH1t)+(TDH2t-ARO3-TEF1p) XII-4::(TDH3p-At4CL-ADH1t) + (TDH2t-RsCHS-CCW12p)+(tHXT7p-PsCHI-FBAt) XII-1::(TDH3p-At4CL-ADH1t) XII-5::(CYC1t-At4CL-TPIp) XI-1::(CYC1t-At4CL-TPIp)* | NAG3-1 | This work |
| NAG4-2 | *MATa ura3-52 can1Δ::cas9-natNT2 TRP1 LEU2 HIS3 XII-2::(GPM1p-AtPAL2-FBA1t)+(TDH3p-AtC4H-CYC1t)+(tHXT7p-AtATR2-pYX212t)+(PGK1p-CYB5-ADH1t) X-3::(TPI1p-EcaroL-pYX212t)+(ADH1t-ARO7* $^{G141S}$*-TEF1p)+(PGK1p-ARO4* $^{K229L}$*-CYC1t) X-4::(CYC1t-ARO1-TPI1p)+(TDH3p-ARO2-ADH1t)+(TDH2t-ARO3-TEF1p) XII-4::(TDH3p-At4CL-ADH1t) + (TDH2t-RsCHS-CCW12p)+(tHXT7p-PsCHI-FBAt) XII-1::(TDH3p-At4CL-ADH1t) + (TDH2t-RsCHS-CCW12p)+(tHXT7p-PsCHI-FBAt) XII-5::(CYC1t-At4CL-TPIp) XI-1::(CYC1t-At4CL-TPIp)* | NAG3-2 | This work |
| NAG4-3 | *MATa ura3-52 can1Δ::cas9-natNT2 TRP1 LEU2 HIS3 XII-2::(GPM1p-AtPAL2-FBA1t)+(TDH3p-AtC4H-CYC1t)+(tHXT7p-AtATR2-pYX212t)+(PGK1p-CYB5-ADH1t) X-3::(TPI1p-EcaroL-pYX212t)+(ADH1t-ARO7* $^{G141S}$*-TEF1p)+(PGK1p-ARO4* $^{K229L}$*-CYC1t) X-4::(CYC1t-ARO1-TPI1p)+(TDH3p-ARO2-ADH1t)+(TDH2t-ARO3-TEF1p) XII-4::(TDH3p-At4CL-ADH1t) + (TDH2t-RsCHS-CCW12p)+(tHXT7p-PsCHI-FBAt) XII-1::(TDH3p-At4CL-ADH1t) + (TDH2t-RsCHS-CCW12p)+(tHXT7p-PsCHI-FBAt) XII-5::(pYX212t-PsCHI-PGKp)+(TEF1p-RsCHS-FBAt) + (CYC1t-At4CL-TPIp) XI-1::(CYC1t-At4CL-TPIp)* | NAG3-3 | This work |
| NAG4-4 | *MATa ura3-52 can1Δ::cas9-natNT2 TRP1 LEU2 HIS3 XII-2::(GPM1p-AtPAL2-FBA1t)+(TDH3p-AtC4H-CYC1t)+(tHXT7p-AtATR2-pYX212t)+(PGK1p-CYB5-ADH1t) X-3::(TPI1p-EcaroL-pYX212t)+(ADH1t-ARO7* $^{G141S}$*-TEF1p)+(PGK1p-ARO4* $^{K229L}$*-CYC1t) X-4::(CYC1t-ARO1-TPI1p)+(TDH3p-ARO2-ADH1t)+(TDH2t-ARO3-TEF1p) XII-4::(TDH3p-At4CL-ADH1t) + (TDH2t-RsCHS-CCW12p)+(tHXT7p-PsCHI-FBAt) XII-1::(TDH3p-At4CL-ADH1t) + (TDH2t-RsCHS-CCW12p)+(tHXT7p-PsCHI-FBAt) XII-5::(pYX212t-PsCHI-PGKp)+(TEF1p-RsCHS-FBAt) + (CYC1t-At4CL-TPIp) XI-1::(pYX212t-PsCHI-PGKp)+(TEF1p-RsCHS-FBAt) + (CYC1t-At4CL-TPIp)* | NAG3-3 | This work |
| NAG-CHS*2 | *MATa ura3-52 can1Δ::cas9-natNT2 TRP1 LEU2 HIS3 XII-2::(GPM1p-AtPAL2-FBA1t)+(TDH3p-AtC4H-CYC1t)+(tHXT7p-AtATR2-pYX212t)+(PGK1p-CYB5-ADH1t) X-3::(TPI1p-EcaroL-pYX212t)+(ADH1t-ARO7* $^{G141S}$*-TEF1p)+(PGK1p-ARO4* $^{K229L}$*-CYC1t) X-4::(CYC1t-ARO1-TPI1p)+(TDH3p-ARO2-ADH1t)+(TDH2t-ARO3-TEF1p) XII-4::(TDH3p-At4CL-ADH1t) + (TDH2t-RsCHS-CCW12p)+(tHXT7p-PsCHI-FBAt) XII-5::(TEF1p-RsCHS-FBAt)* | NAG10 | This work |
| NAG-CHS*3 | *MATa ura3-52 can1Δ::cas9-natNT2 TRP1 LEU2 HIS3 XII-2::(GPM1p-AtPAL2-FBA1t)+(TDH3p-AtC4H-CYC1t)+(tHXT7p-AtATR2-pYX212t)+(PGK1p-CYB5-ADH1t) X-3::(TPI1p-EcaroL-pYX212t)+(ADH1t-ARO7* $^{G141S}$*-TEF1p)+(PGK1p-ARO4* $^{K229L}$*-CYC1t) X-4::(CYC1t-ARO1-TPI1p)+(TDH3p-ARO2-ADH1t)+(TDH2t-ARO3-TEF1p) XII-4::(TDH3p-At4CL-ADH1t) + (TDH2t-RsCHS-CCW12p)+(tHXT7p-PsCHI-FBAt) XII-5::(TEF1p-RsCHS-FBAt) XI-1: (TEF1p-RsCHS-FBAt)* | NAG-CHS*2 | This work |

| | | | |
|---|---|---|---|
| NAG-CHI*2 | *MATa ura3-52 can1Δ::cas9-natNT2 TRP1 LEU2 HIS3 XII-2::(GPM1p-AtPAL2-FBA1t)+(TDH3p-AtC4H-CYC1t)+(tHXT7p-AtATR2-pYX212t)+(PGK1p-CYB5-ADH1t) X-3::(TPI1p-EcaroL-pYX212t)+(ADH1t-ARO7*$^{G141S}$*-TEF1p)+(PGK1p-ARO4*$^{K229L}$*-CYC1t) X-4::(CYC1t-ARO1-TPI1p)+(TDH3p-ARO2-ADH1t)+(TDH2t-ARO3-TEF1p) XII-4::(TDH3p-At4CL-ADH1t) + (TDH2t-RsCHS-CCW12p)+(tHXT7p-PsCHI-FBAt) XII-5::(pYX212t-PsCHI-PGKp)* | NAG10 | This work |
| NAG-CHI*3 | *MATa ura3-52 can1Δ::cas9-natNT2 TRP1 LEU2 HIS3 XII-2::(GPM1p-AtPAL2-FBA1t)+(TDH3p-AtC4H-CYC1t)+(tHXT7p-AtATR2-pYX212t)+(PGK1p-CYB5-ADH1t) X-3::(TPI1p-EcaroL-pYX212t)+(ADH1t-ARO7*$^{G141S}$*-TEF1p)+(PGK1p-ARO4*$^{K229L}$*-CYC1t) X-4::(CYC1t-ARO1-TPI1p)+(TDH3p-ARO2-ADH1t)+(TDH2t-ARO3-TEF1p) XII-4::(TDH3p-At4CL-ADH1t) + (TDH2t-RsCHS-CCW12p)+(tHXT7p-PsCHI-FBAt) XII-5::(pYX212t-PsCHI-PGKp) XI-1:(pYX212t-PsCHI-PGKp)* | NAG-CHI*2 | This work |
| NAG2-5 | *MATa ura3-52 can1Δ::cas9-natNT2 TRP1 LEU2 HIS3 XII-2::(GPM1p-AtPAL2-FBA1t)+(TDH3p-AtC4H-CYC1t)+(tHXT7p-AtATR2-pYX212t)+(PGK1p-CYB5-ADH1t) X-3::(TPI1p-EcaroL-pYX212t)+(ADH1t-ARO7*$^{G141S}$*-TEF1p)+(PGK1p-ARO4*$^{K229L}$*-CYC1t) X-4::(CYC1t-ARO1-TPI1p)+(TDH3p-ARO2-ADH1t)+(TDH2t-ARO3-TEF1p) XII-4::(TDH3p-At4CL-ADH1t) + (TDH2t-RsCHS-CCW12p)+(tHXT7p-PsCHI-FBAt) XII-1::(TDH3p-At4CL-ADH1t) + (TDH2t-RsCHS-CCW12p)+(tHXT7p-PsCHI-FBAt) XII-5::(pYX212t-PsCHI-PGKp)+ (TEF1p-RsCHS-FBAt) XI-1::(pYX212t-PsCHI-PGKp)+ (TEF1p-RsCHS-FBAt) XI-3::(TDH2t-RsCHS-CCW12p)+(tHXT7p-PsCHI-FBAt)* | NAG2-4 | This work |
| NAG3-5 | *MATa ura3-52 can1Δ::cas9-natNT2 TRP1 LEU2 HIS3 XII-2::(GPM1p-AtPAL2-FBA1t)+(TDH3p-AtC4H-CYC1t)+(tHXT7p-AtATR2-pYX212t)+(PGK1p-CYB5-ADH1t) X-3::(TPI1p-EcaroL-pYX212t)+(ADH1t-ARO7*$^{G141S}$*-TEF1p)+(PGK1p-ARO4*$^{K229L}$*-CYC1t) X-4::(CYC1t-ARO1-TPI1p)+(TDH3p-ARO2-ADH1t)+(TDH2t-ARO3-TEF1p) XII-4::(TDH3p-At4CL-ADH1t) + (TDH2t-RsCHS-CCW12p)+(tHXT7p-PsCHI-FBAt) XII-1::(TDH3p-At4CL-ADH1t) + (TDH2t-RsCHS-CCW12p)+(tHXT7p-PsCHI-FBAt) XII-5::(pYX212t-PsCHI-PGKp)+ (TEF1p-RsCHS-FBAt) + (CYC1t-At4CL-TPIp) XI-1::(pYX212t-PsCHI-PGKp)+ (TEF1p-RsCHS-FBAt) XI-3::(TDH2t-RsCHS-CCW12p)+(tHXT7p-PsCHI-FBAt)* | NAG3-4 | This work |
| NAG4-5 | *MATa ura3-52 can1Δ::cas9-natNT2 TRP1 LEU2 HIS3 XII-2::(GPM1p-AtPAL2-FBA1t)+(TDH3p-AtC4H-CYC1t)+(tHXT7p-AtATR2-pYX212t)+(PGK1p-CYB5-ADH1t) X-3::(TPI1p-EcaroL-pYX212t)+(ADH1t-ARO7*$^{G141S}$*-TEF1p)+(PGK1p-ARO4*$^{K229L}$*-CYC1t) X-4::(CYC1t-ARO1-TPI1p)+(TDH3p-ARO2-ADH1t)+(TDH2t-ARO3-TEF1p) XII-4::(TDH3p-At4CL-ADH1t) + (TDH2t-RsCHS-CCW12p)+(tHXT7p-PsCHI-FBAt) XII-1::(TDH3p-At4CL-ADH1t) + (TDH2t-RsCHS-CCW12p)+(tHXT7p-PsCHI-FBAt) XII-5::(pYX212t-PsCHI-PGKp)+ (TEF1p-RsCHS-FBAt) + (CYC1t-At4CL-TPIp) XI-1::(pYX212t-PsCHI-PGKp)+ (TEF1p-RsCHS-FBAt) + (CYC1t-At4CL-TPIp) XI-3:;(TDH2t-RsCHS-CCW12p)+(tHXT7p-PsCHI-FBAt)* | NAG4-4 | This work |
| NAG12 | *MATa ura3-52 can1Δ::cas9-natNT2 TRP1 LEU2 HIS3 XII-2::(GPM1p-AtPAL2-FBA1t)+(TDH3p-AtC4H-CYC1t)+(tHXT7p-AtATR2-pYX212t)+(PGK1p-CYB5-ADH1t) X-3::(TPI1p-EcaroL-pYX212t)+(ADH1t-ARO7*$^{G141S}$*-TEF1p)+(PGK1p-ARO4*$^{K229L}$*-CYC1t) X-4::(CYC1t-ARO1-TPI1p)+(TDH3p-ARO2-ADH1t)+(TDH2t-ARO3-TEF1p) XII-4::(TDH3p-At4CL-ADH1t) + (TDH2t-RsCHS-CCW12p)+(tHXT7p-PsCHI-FBAt) XI-2::(TPIp-ACC1*$^{S659A,S1157A}$* -TDH2t)* | NAG010 | This work |
| NAG13 | *MATa ura3-52 can1Δ::cas9-natNT2 TRP1 LEU2 HIS3 XII-2::(GPM1p-AtPAL2-FBA1t)+(TDH3p-AtC4H-CYC1t)+(tHXT7p-AtATR2-pYX212t)+(PGK1p-CYB5-ADH1t) X-3::(TPI1p-EcaroL-pYX212t)+(ADH1t-ARO7*$^{G141S}$*-TEF1p)+(PGK1p-ARO4*$^{K229L}$*-CYC1t) X-4::(CYC1t-ARO1-TPI1p)+(TDH3p-ARO2-ADH1t)+(TDH2t-ARO3-TEF1p) XII-4::(TDH3p-At4CL-ADH1t) + (TDH2t-RsCHS-CCW12p)+(tHXT7p-PsCHI-FBAt) XII-3::(TDH2t-matB-TDH3p) + (tHXT7p-matC-CYC1t)* | NAG010 | This work |
| NAG14 | *MATa ura3-52 can1Δ::cas9-natNT2 TRP1 LEU2 HIS3 XII-2::(GPM1p-AtPAL2-FBA1t)+(TDH3p-AtC4H-CYC1t)+(tHXT7p-AtATR2-pYX212t)+(PGK1p-CYB5-ADH1t) X-3::(TPI1p-EcaroL-pYX212t)+(ADH1t-ARO7*$^{G141S}$*-TEF1p)+(PGK1p-ARO4*$^{K229L}$*-CYC1t) X-4::(CYC1t-ARO1-TPI1p)+(TDH3p-ARO2-ADH1t)+(TDH2t-ARO3-TEF1p) XII-4::(TDH3p-At4CL-ADH1t) + (TDH2t-RsCHS-CCW12p)+(tHXT7p-PsCHI-FBAt) XI-2::(TPIp-ACC1*$^{S659A,S1157A}$* -TDH2t) XII-3::(TDH2t-matB-TDH3p) + (tHXT7p-matC-CYC1t)* | NAG12 | This work |

| | | | |
|---|---|---|---|
| NAG37 | *MATa ura3-52 can1Δ::cas9-natNT2 TRP1 LEU2 HIS3 XII-2::(GPM1p-AtPAL2-FBA1t)+(TDH3p-AtC4H-CYC1t)+(tHXT7p-AtATR2-pYX212t)+(PGK1p-CYB5-ADH1t) X-3::(TPI1p-EcaroL-pYX212t)+(ADH1t-ARO7 $^{G141S}$-TEF1p)+(PGK1p-ARO4 $^{K229L}$-CYC1t) X-4::(CYC1t-ARO1-TPI1p)+(TDH3p-ARO2-ADH1t)+(TDH2t-ARO3-TEF1p) XII-4::(TDH3p-At4CL-ADH1t) + (TDH2t-RsCHS-CCW12p)+(tHXT7p-PsCHI-FBAt) XII-1::(TDH3p-At4CL-ADH1t) + (TDH2t-RsCHS-CCW12p)+(tHXT7p-PsCHI-FBAt) XII-5::(pYX212t-PsCHI-PGKp)+ (TEF1p-RsCHS-FBAt) + (CYC1t-At4CL-TPIp) XI-1::(pYX212t-PsCHI-PGKp)+ (TEF1p-RsCHS-FBAt) XI-2: (TPIp-ACC1 $^{S659A,S1157A}$ -TDH2t)* | NAG3-4 | This work |
| NAG38 | *MATa ura3-52 can1Δ::cas9-natNT2 TRP1 LEU2 HIS3 XII-2::(GPM1p-AtPAL2-FBA1t)+(TDH3p-AtC4H-CYC1t)+(tHXT7p-AtATR2-pYX212t)+(PGK1p-CYB5-ADH1t) X-3::(TPI1p-EcaroL-pYX212t)+(ADH1t-ARO7 $^{G141S}$-TEF1p)+(PGK1p-ARO4 $^{K229L}$-CYC1t) X-4::(CYC1t-ARO1-TPI1p)+(TDH3p-ARO2-ADH1t)+(TDH2t-ARO3-TEF1p) XII-4::(TDH3p-At4CL-ADH1t) + (TDH2t-RsCHS-CCW12p)+(tHXT7p-PsCHI-FBAt) XII-1::(TDH3p-At4CL-ADH1t) + (TDH2t-RsCHS-CCW12p)+(tHXT7p-PsCHI-FBAt) XII-5::(pYX212t-PsCHI-PGKp)+ (TEF1p-RsCHS-FBAt) + (CYC1t-At4CL-TPIp) XI-1::(pYX212t-PsCHI-PGKp)+ (TEF1p-RsCHS-FBAt) XII-3::(TDH2t-matB-TDH3p) + (tHXT7p-matC-CYC1t)* | NAG3-4 | This work |
| NAG39 | *MATa ura3-52 can1Δ::cas9-natNT2 TRP1 LEU2 HIS3 XII-2::(GPM1p-AtPAL2-FBA1t)+(TDH3p-AtC4H-CYC1t)+(tHXT7p-AtATR2-pYX212t)+(PGK1p-CYB5-ADH1t) X-3::(TPI1p-EcaroL-pYX212t)+(ADH1t-ARO7 $^{G141S}$-TEF1p)+(PGK1p-ARO4 $^{K229L}$-CYC1t) X-4::(CYC1t-ARO1-TPI1p)+(TDH3p-ARO2-ADH1t)+(TDH2t-ARO3-TEF1p) XII-4::(TDH3p-At4CL-ADH1t) + (TDH2t-RsCHS-CCW12p)+(tHXT7p-PsCHI-FBAt) XII-1::(TDH3p-At4CL-ADH1t) + (TDH2t-RsCHS-CCW12p)+(tHXT7p-PsCHI-FBAt) XII-5::(pYX212t-PsCHI-PGKp)+ (TEF1p-RsCHS-FBAt) + (CYC1t-At4CL-TPIp) XI-1::(pYX212t-PsCHI-PGKp)+ (TEF1p-RsCHS-FBAt) XI-2::(TPIp-ACC1 $^{S659A,S1157A}$ -TDH2t) XII-3::(TDH2t-matB-TDH3p) + (tHXT7p-matC-CYC1t)* | NAG37 | This work |
| NAG41 | *MATa ura3-52 can1Δ::cas9-natNT2 TRP1 LEU2 HIS3 X-3::(TPI1p-EcaroL-pYX212t)+(ADH1t-ARO7 $^{G141S}$-TEF1p)+(PGK1p-ARO4 $^{K229L}$-CYC1t) X-4::(CYC1t-ARO1-TPI1p)+(TDH3p-ARO2-ADH1t)+(TDH2t-ARO3-TEF1p) XII-4::(TDH3p-At4CL-ADH1t) + (TDH2t-RsCHS-CCW12p)+(tHXT7p-PsCHI-FBAt) XII-1::(TDH3p-At4CL-ADH1t) + (TDH2t-RsCHS-CCW12p)+(tHXT7p-PsCHI-FBAt) XII-5::(pYX212t-PsCHI-PGKp)+ (TEF1p-RsCHS-FBAt) + (CYC1t-At4CL-TPIp) XI-1::(pYX212t-PsCHI-PGKp)+ (TEF1p-RsCHS-FBAt) XI-2::(TPIp-ACC1 $^{S659A,S1157A}$ -TDH2t) XII-3::(TDH2t-matB-TDH3p) + (tHXT7p-matC-CYC1t)* | NAG39 | This work |
| NAG58 | *MATa ura3-52 can1Δ::cas9-natNT2 TRP1 LEU2 HIS3 XII-2:: TDH3p-AtC4H-CYC1t)+(tHXT7p-AtATR2-pYX212t)+(PGK1p-CYB5-ADH1t) X-3::(TPI1p-EcaroL-pYX212t)+(ADH1t-ARO7 $^{G141S}$-TEF1p)+(PGK1p-ARO4 $^{K229L}$-CYC1t) X-4::(CYC1t-ARO1-TPI1p)+(TDH3p-ARO2-ADH1t)+(TDH2t-ARO3-TEF1p) XII-4::(TDH3p-At4CL-ADH1t) + (TDH2t-RsCHS-CCW12p)+(tHXT7p-PsCHI-FBAt) XII-1::(TDH3p-At4CL-ADH1t) + (TDH2t-RsCHS-CCW12p)+(tHXT7p-PsCHI-FBAt) XII-5::(pYX212t-PsCHI-PGKp)+ (TEF1p-RsCHS-FBAt) + (CYC1t-At4CL-TPIp) XI-1::(pYX212t-PsCHI-PGKp)+ (TEF1p-RsCHS-FBAt) XI-2::(TPIp-ACC1 $^{S659A,S1157A}$ -TDH2t) XII-3::(TDH2t-matB-TDH3p) + (tHXT7p-matC-CYC1t) XI-3:: (GPM1p-AtPAL2-FBA1t)* | NAG39 | This work |
| NAG59 | *MATa ura3-52 can1Δ::cas9-natNT2 TRP1 LEU2 HIS3 XII-2:: TDH3p-AtC4H-CYC1t)+(tHXT7p-AtATR2-pYX212t)+(PGK1p-CYB5-ADH1t) X-3::(TPI1p-EcaroL-pYX212t)+(ADH1t-ARO7 $^{G141S}$-TEF1p)+(PGK1p-ARO4 $^{K229L}$-CYC1t) X-4::(CYC1t-ARO1-TPI1p)+(TDH3p-ARO2-ADH1t)+(TDH2t-ARO3-TEF1p) XII-4::(TDH3p-At4CL-ADH1t) + (TDH2t-RsCHS-CCW12p)+(tHXT7p-PsCHI-FBAt) XII-1::(TDH3p-At4CL-ADH1t) + (TDH2t-RsCHS-CCW12p)+(tHXT7p-PsCHI-FBAt) XII-5::(pYX212t-PsCHI-PGKp)+ (TEF1p-RsCHS-FBAt) + (CYC1t-At4CL-TPIp) XI-1::(pYX212t-PsCHI-PGKp)+ (TEF1p-RsCHS-FBAt) XI-2::(TPIp-ACC1 $^{S659A,S1157A}$ -TDH2t) XII-3::(TDH2t-matB-TDH3p) + (tHXT7p-matC-CYC1t) XI-3:: (CCW12p-AtPAL2-FBA1t)* | NAG39 | This work |
| NAG60 | *MATa ura3-52 can1Δ::cas9-natNT2 TRP1 LEU2 HIS3 XII-2:: TDH3p-AtC4H-CYC1t)+(tHXT7p-AtATR2-pYX212t)+(PGK1p-CYB5-ADH1t) X-3::(TPI1p-EcaroL-pYX212t)+(ADH1t-ARO7 $^{G141S}$-TEF1p)+(PGK1p-ARO4 $^{K229L}$-CYC1t) X-4::(CYC1t-ARO1-TPI1p)+(TDH3p-ARO2-ADH1t)+(TDH2t-ARO3-TEF1p) XII-4::(TDH3p-At4CL-ADH1t) + (TDH2t-RsCHS-CCW12p)+(tHXT7p-PsCHI-FBAt) XII-1::(TDH3p-At4CL-ADH1t) + (TDH2t-RsCHS-CCW12p)+(tHXT7p-PsCHI-* | NAG39 | This work |

| | | | |
|---|---|---|---|
| | *FBAt) XII-5::(pYX212t-PsCHI-PGKp)+ (TEF1p-RsCHS-FBAt) + (CYC1t-At4CL-TPIp) XI-1::(pYX212t-PsCHI-PGKp)+ (TEF1p-RsCHS-FBAt) XI-2::(TPIp-ACC1*$^{S659A,S1157A}$ *-TDH2t) XII-3::(TDH2t-matB-TDH3p) + (tHXT7p-matC-CYC1t) XI-3:: (CCW12_BS2p-AtPAL2-FBA1t)* | | |
| NAG61 | *MATa ura3-52 can1Δ::cas9-natNT2 TRP1 LEU2 HIS3 XII-2:: TDH3p-AtC4H-CYC1t)+(tHXT7p-AtATR2-pYX212t)+(PGK1p-CYB5-ADH1t) X-3::(TPI1p-EcaroL-pYX212t)+(ADH1t-ARO7*$^{G141S}$*-TEF1p)+(PGK1p-ARO4*$^{K229L}$*-CYC1t) X-4::(CYC1t-ARO1-TPI1p)+(TDH3p-ARO2-ADH1t)+(TDH2t-ARO3-TEF1p) XII-4::(TDH3p-At4CL-ADH1t) + (TDH2t-RsCHS-CCW12p)+(tHXT7p-PsCHI-FBAt) XII-1::(TDH3p-At4CL-ADH1t) + (TDH2t-RsCHS-CCW12p)+(tHXT7p-PsCHI-FBAt) XII-5::(pYX212t-PsCHI-PGKp)+ (TEF1p-RsCHS-FBAt) + (CYC1t-At4CL-TPIp) XI-1::(pYX212t-PsCHI-PGKp)+ (TEF1p-RsCHS-FBAt) XI-2::(TPIp-ACC1*$^{S659A,S1157A}$ *-TDH2t) XII-3::(TDH2t-matB-TDH3p) + (tHXT7p-matC-CYC1t) XI-3:: (CCW12_BS123p-AtPAL2-FBA1t)* | NAG39 | This work |
| NAG62 | *MATa ura3-52 can1Δ::cas9-natNT2 TRP1 LEU2 HIS3 XII-2:: TDH3p-AtC4H-CYC1t)+(tHXT7p-AtATR2-pYX212t)+(PGK1p-CYB5-ADH1t) X-3::(TPI1p-EcaroL-pYX212t)+(ADH1t-ARO7*$^{G141S}$*-TEF1p)+(PGK1p-ARO4*$^{K229L}$*-CYC1t) X-4::(CYC1t-ARO1-TPI1p)+(TDH3p-ARO2-ADH1t)+(TDH2t-ARO3-TEF1p) XII-4::(TDH3p-At4CL-ADH1t) + (TDH2t-RsCHS-CCW12p)+(tHXT7p-PsCHI-FBAt) XII-1::(TDH3p-At4CL-ADH1t) + (TDH2t-RsCHS-CCW12p)+(tHXT7p-PsCHI-FBAt) XII-5::(pYX212t-PsCHI-PGKp)+ (TEF1p-RsCHS-FBAt) + (CYC1t-At4CL-TPIp) XI-1::(pYX212t-PsCHI-PGKp)+ (TEF1p-RsCHS-FBAt) XI-2::(TPIp-ACC1*$^{S659A,S1157A}$ *-TDH2t) XII-3::(TDH2t-matB-TDH3p) + (tHXT7p-matC-CYC1t) XI-3:: (TDH3p-AtPAL2-FBA1t)* | NAG39 | This work |
| NAG63 | *MATa ura3-52 can1Δ::cas9-natNT2 TRP1 LEU2 HIS3 XII-2:: TDH3p-AtC4H-CYC1t)+(tHXT7p-AtATR2-pYX212t)+(PGK1p-CYB5-ADH1t) X-3::(TPI1p-EcaroL-pYX212t)+(ADH1t-ARO7*$^{G141S}$*-TEF1p)+(PGK1p-ARO4*$^{K229L}$*-CYC1t) X-4::(CYC1t-ARO1-TPI1p)+(TDH3p-ARO2-ADH1t)+(TDH2t-ARO3-TEF1p) XII-4::(TDH3p-At4CL-ADH1t) + (TDH2t-RsCHS-CCW12p)+(tHXT7p-PsCHI-FBAt) XII-1::(TDH3p-At4CL-ADH1t) + (TDH2t-RsCHS-CCW12p)+(tHXT7p-PsCHI-FBAt) XII-5::(pYX212t-PsCHI-PGKp)+ (TEF1p-RsCHS-FBAt) + (CYC1t-At4CL-TPIp) XI-1::(pYX212t-PsCHI-PGKp)+ (TEF1p-RsCHS-FBAt) XI-2::(TPIp-ACC1*$^{S659A,S1157A}$ *-TDH2t) XII-3::(TDH2t-matB-TDH3p) + (tHXT7p-matC-CYC1t) XI-3:: (TDH3_BS23p-AtPAL2-FBA1t)* | NAG39 | This work |
| NAG64 | *MATa ura3-52 can1Δ::cas9-natNT2 TRP1 LEU2 HIS3 XII-2:: TDH3p-AtC4H-CYC1t)+(tHXT7p-AtATR2-pYX212t)+(PGK1p-CYB5-ADH1t) X-3::(TPI1p-EcaroL-pYX212t)+(ADH1t-ARO7*$^{G141S}$*-TEF1p)+(PGK1p-ARO4*$^{K229L}$*-CYC1t) X-4::(CYC1t-ARO1-TPI1p)+(TDH3p-ARO2-ADH1t)+(TDH2t-ARO3-TEF1p) XII-4::(TDH3p-At4CL-ADH1t) + (TDH2t-RsCHS-CCW12p)+(tHXT7p-PsCHI-FBAt) XII-1::(TDH3p-At4CL-ADH1t) + (TDH2t-RsCHS-CCW12p)+(tHXT7p-PsCHI-FBAt) XII-5::(pYX212t-PsCHI-PGKp)+ (TEF1p-RsCHS-FBAt) + (CYC1t-At4CL-TPIp) XI-1::(pYX212t-PsCHI-PGKp)+ (TEF1p-RsCHS-FBAt) XI-2::(TPIp-ACC1*$^{S659A,S1157A}$ *-TDH2t) XII-3::(TDH2t-matB-TDH3p) + (tHXT7p-matC-CYC1t) XI-3:: (TEF1p-AtPAL2-FBA1t)* | NAG39 | This work |
| NAG65 | *MATa ura3-52 can1Δ::cas9-natNT2 TRP1 LEU2 HIS3 XII-2:: TDH3p-AtC4H-CYC1t)+(tHXT7p-AtATR2-pYX212t)+(PGK1p-CYB5-ADH1t) X-3::(TPI1p-EcaroL-pYX212t)+(ADH1t-ARO7*$^{G141S}$*-TEF1p)+(PGK1p-ARO4*$^{K229L}$*-CYC1t) X-4::(CYC1t-ARO1-TPI1p)+(TDH3p-ARO2-ADH1t)+(TDH2t-ARO3-TEF1p) XII-4::(TDH3p-At4CL-ADH1t) + (TDH2t-RsCHS-CCW12p)+(tHXT7p-PsCHI-FBAt) XII-1::(TDH3p-At4CL-ADH1t) + (TDH2t-RsCHS-CCW12p)+(tHXT7p-PsCHI-FBAt) XII-5::(pYX212t-PsCHI-PGKp)+ (TEF1p-RsCHS-FBAt) + (CYC1t-At4CL-TPIp) XI-1::(pYX212t-PsCHI-PGKp)+ (TEF1p-RsCHS-FBAt) XI-2::(TPIp-ACC1*$^{S659A,S1157A}$ *-TDH2t) XII-3::(TDH2t-matB-TDH3p) + (tHXT7p-matC-CYC1t) XI-3:: (TEF1_BS123p-AtPAL2-FBA1t)* | NAG39 | This work |
| NAG39-2 | *MATa ura3-52 can1Δ::cas9-natNT2 TRP1 LEU2 HIS3 XII-2::(GPM1p-AtPAL2-FBA1t)+(TDH3p-AtC4H-CYC1t)+(tHXT7p-AtATR2-pYX212t)+(PGK1p-CYB5-ADH1t) X-3::(TPI1p-EcaroL-pYX212t)+(ADH1t-ARO7*$^{G141S}$*-TEF1p)+(PGK1p-ARO4*$^{K229L}$*-CYC1t) X-4::(CYC1t-ARO1-TPI1p)+(TDH3p-ARO2-ADH1t)+(TDH2t-ARO3-TEF1p) XII-4::(TDH3p-At4CL-ADH1t) + (TDH2t-RsCHS-CCW12p)+(tHXT7p-PsCHI-FBAt) XII-1::(TDH3p-At4CL-ADH1t) + (TDH2t-RsCHS-CCW12p)+(tHXT7p-PsCHI-FBAt) XII-5::(pYX212t-PsCHI-PGKp)+ (TEF1p-RsCHS-FBAt) + (CYC1t-At4CL-TPIp) XI-1::(pYX212t-PsCHI-PGKp)+* | NAG39 | This work |

| | | | |
|---|---|---|---|
| | *(TEF1p-RsCHS-FBAt) XI-2::(TPIp-ACC1* $^{S659A,S1157A}$ *-TDH2t) XII-3::(TDH2t-matB-TDH3p) + (tHXT7p-matC-CYC1t) X-2::(TEF1-NLS_FapR-ADH1t)* | | |
| NAG66 | *MATa ura3-52 can1Δ::cas9-natNT2 TRP1 LEU2 HIS3 XII-2:: TDH3p-AtC4H-CYC1t)+(tHXT7p-AtATR2-pYX212t)+(PGK1p-CYB5-ADH1t) X-3::(TPI1p-EcaroL-pYX212t)+(ADH1t-ARO7* $^{G141S}$*-TEF1p)+(PGK1p-ARO4* $^{K229L}$*-CYC1t) X-4::(CYC1t-ARO1-TPI1p)+(TDH3p-ARO2-ADH1t)+(TDH2t-ARO3-TEF1p) XII-4::(TDH3p-At4CL-ADH1t) + (TDH2t-RsCHS-CCW12p)+(tHXT7p-PsCHI-FBAt) XII-1::(TDH3p-At4CL-ADH1t) + (TDH2t-RsCHS-CCW12p)+(tHXT7p-PsCHI-FBAt) XII-5::(pYX212t-PsCHI-PGKp)+ (TEF1p-RsCHS-FBAt) + (CYC1t-At4CL-TPIp) XI-1::(pYX212t-PsCHI-PGKp)+ (TEF1p-RsCHS-FBAt) XI-2::(TPIp-ACC1* $^{S659A,S1157A}$ *-TDH2t) XII-3::(TDH2t-matB-TDH3p) + (tHXT7p-matC-CYC1t) XI-3:: (GPM1p-AtPAL2-FBA1t) X-2::(TEF1-NLS_FapR-ADH1t)* | NAG58 | This work |
| NAG67 | *MATa ura3-52 can1Δ::cas9-natNT2 TRP1 LEU2 HIS3 XII-2:: TDH3p-AtC4H-CYC1t)+(tHXT7p-AtATR2-pYX212t)+(PGK1p-CYB5-ADH1t) X-3::(TPI1p-EcaroL-pYX212t)+(ADH1t-ARO7* $^{G141S}$*-TEF1p)+(PGK1p-ARO4* $^{K229L}$*-CYC1t) X-4::(CYC1t-ARO1-TPI1p)+(TDH3p-ARO2-ADH1t)+(TDH2t-ARO3-TEF1p) XII-4::(TDH3p-At4CL-ADH1t) + (TDH2t-RsCHS-CCW12p)+(tHXT7p-PsCHI-FBAt) XII-1::(TDH3p-At4CL-ADH1t) + (TDH2t-RsCHS-CCW12p)+(tHXT7p-PsCHI-FBAt) XII-5::(pYX212t-PsCHI-PGKp)+ (TEF1p-RsCHS-FBAt) + (CYC1t-At4CL-TPIp) XI-1::(pYX212t-PsCHI-PGKp)+ (TEF1p-RsCHS-FBAt) XI-2::(TPIp-ACC1* $^{S659A,S1157A}$ *-TDH2t) XII-3::(TDH2t-matB-TDH3p) + (tHXT7p-matC-CYC1t) XI-3:: (CCW12p-AtPAL2-FBA1t) X-2::(TEF1-NLS_FapR-ADH1t)* | NAG59 | This work |
| NAG68 | *MATa ura3-52 can1Δ::cas9-natNT2 TRP1 LEU2 HIS3 XII-2:: TDH3p-AtC4H-CYC1t)+(tHXT7p-AtATR2-pYX212t)+(PGK1p-CYB5-ADH1t) X-3::(TPI1p-EcaroL-pYX212t)+(ADH1t-ARO7* $^{G141S}$*-TEF1p)+(PGK1p-ARO4* $^{K229L}$*-CYC1t) X-4::(CYC1t-ARO1-TPI1p)+(TDH3p-ARO2-ADH1t)+(TDH2t-ARO3-TEF1p) XII-4::(TDH3p-At4CL-ADH1t) + (TDH2t-RsCHS-CCW12p)+(tHXT7p-PsCHI-FBAt) XII-1::(TDH3p-At4CL-ADH1t) + (TDH2t-RsCHS-CCW12p)+(tHXT7p-PsCHI-FBAt) XII-5::(pYX212t-PsCHI-PGKp)+ (TEF1p-RsCHS-FBAt) + (CYC1t-At4CL-TPIp) XI-1::(pYX212t-PsCHI-PGKp)+ (TEF1p-RsCHS-FBAt) XI-2::(TPIp-ACC1* $^{S659A,S1157A}$ *-TDH2t) XII-3::(TDH2t-matB-TDH3p) + (tHXT7p-matC-CYC1t) XI-3:: (CCW12_BS2p-AtPAL2-FBA1t) X-2::(TEF1-NLS_FapR-ADH1t)* | NAG60 | This work |
| NAG69 | *MATa ura3-52 can1Δ::cas9-natNT2 TRP1 LEU2 HIS3 XII-2:: TDH3p-AtC4H-CYC1t)+(tHXT7p-AtATR2-pYX212t)+(PGK1p-CYB5-ADH1t) X-3::(TPI1p-EcaroL-pYX212t)+(ADH1t-ARO7* $^{G141S}$*-TEF1p)+(PGK1p-ARO4* $^{K229L}$*-CYC1t) X-4::(CYC1t-ARO1-TPI1p)+(TDH3p-ARO2-ADH1t)+(TDH2t-ARO3-TEF1p) XII-4::(TDH3p-At4CL-ADH1t) + (TDH2t-RsCHS-CCW12p)+(tHXT7p-PsCHI-FBAt) XII-1::(TDH3p-At4CL-ADH1t) + (TDH2t-RsCHS-CCW12p)+(tHXT7p-PsCHI-FBAt) XII-5::(pYX212t-PsCHI-PGKp)+ (TEF1p-RsCHS-FBAt) + (CYC1t-At4CL-TPIp) XI-1::(pYX212t-PsCHI-PGKp)+ (TEF1p-RsCHS-FBAt) XI-2::(TPIp-ACC1* $^{S659A,S1157A}$ *-TDH2t) XII-3::(TDH2t-matB-TDH3p) + (tHXT7p-matC-CYC1t) XI-3:: (CCW12_BS123p-AtPAL2-FBA1t) X-2::(TEF1-NLS_FapR-ADH1t)* | NAG61 | This work |
| NAG70 | *MATa ura3-52 can1Δ::cas9-natNT2 TRP1 LEU2 HIS3 XII-2:: TDH3p-AtC4H-CYC1t)+(tHXT7p-AtATR2-pYX212t)+(PGK1p-CYB5-ADH1t) X-3::(TPI1p-EcaroL-pYX212t)+(ADH1t-ARO7* $^{G141S}$*-TEF1p)+(PGK1p-ARO4* $^{K229L}$*-CYC1t) X-4::(CYC1t-ARO1-TPI1p)+(TDH3p-ARO2-ADH1t)+(TDH2t-ARO3-TEF1p) XII-4::(TDH3p-At4CL-ADH1t) + (TDH2t-RsCHS-CCW12p)+(tHXT7p-PsCHI-FBAt) XII-1::(TDH3p-At4CL-ADH1t) + (TDH2t-RsCHS-CCW12p)+(tHXT7p-PsCHI-FBAt) XII-5::(pYX212t-PsCHI-PGKp)+ (TEF1p-RsCHS-FBAt) + (CYC1t-At4CL-TPIp) XI-1::(pYX212t-PsCHI-PGKp)+ (TEF1p-RsCHS-FBAt) XI-2::(TPIp-ACC1* $^{S659A,S1157A}$ *-TDH2t) XII-3::(TDH2t-matB-TDH3p) + (tHXT7p-matC-CYC1t) XI-3:: (TDH3p-AtPAL2-FBA1t) X-2::(TEF1-NLS_FapR-ADH1t)* | NAG62 | This work |
| NAG71 | *MATa ura3-52 can1Δ::cas9-natNT2 TRP1 LEU2 HIS3 XII-2:: TDH3p-AtC4H-CYC1t)+(tHXT7p-AtATR2-pYX212t)+(PGK1p-CYB5-ADH1t) X-3::(TPI1p-EcaroL-pYX212t)+(ADH1t-ARO7* $^{G141S}$*-TEF1p)+(PGK1p-ARO4* $^{K229L}$*-CYC1t) X-4::(CYC1t-ARO1-TPI1p)+(TDH3p-ARO2-ADH1t)+(TDH2t-ARO3-TEF1p) XII-4::(TDH3p-At4CL-ADH1t) + (TDH2t-RsCHS-CCW12p)+(tHXT7p-PsCHI-FBAt) XII-1::(TDH3p-At4CL-ADH1t) + (TDH2t-RsCHS-CCW12p)+(tHXT7p-PsCHI-FBAt) XII-5::(pYX212t-PsCHI-PGKp)+ (TEF1p-RsCHS-FBAt) + (CYC1t-At4CL-TPIp) XI-1::(pYX212t-PsCHI-PGKp)+ (TEF1p-RsCHS-FBAt) XI-2::(TPIp-ACC1* $^{S659A,S1157A}$ *-TDH2t) XII-3::(TDH2t-matB-TDH3p) + (tHXT7p-matC-CYC1t) XI-3:: (TDH3_BS23p-AtPAL2-FBA1t) X-2::(TEF1-NLS_FapR-ADH1t)* | NAG63 | This work |

| | | | |
|---|---|---|---|
| NAG72 | *MATa ura3-52 can1Δ::cas9-natNT2 TRP1 LEU2 HIS3 XII-2:: TDH3p-AtC4H-CYC1t)+(tHXT7p-AtATR2-pYX212t)+(PGK1p-CYB5-ADH1t) X-3::(TPI1p-EcaroL-pYX212t)+(ADH1t-ARO7* $^{G141S}$*-TEF1p)+(PGK1p-ARO4* $^{K229L}$*-CYC1t) X-4::(CYC1t-ARO1-TPI1p)+(TDH3p-ARO2-ADH1t)+(TDH2t-ARO3-TEF1p) XII-4::(TDH3p-At4CL-ADH1t) + (TDH2t-RsCHS-CCW12p)+(tHXT7p-PsCHI-FBAt) XII-1::(TDH3p-At4CL-ADH1t) + (TDH2t-RsCHS-CCW12p)+(tHXT7p-PsCHI-FBAt) XII-5::(pYX212t-PsCHI-PGKp)+ (TEF1p-RsCHS-FBAt) + (CYC1t-At4CL-TPIp) XI-1::(pYX212t-PsCHI-PGKp)+ (TEF1p-RsCHS-FBAt) XI-2::(TPIp-ACC1* $^{S659A,S1157A}$ *-TDH2t) XII-3::(TDH2t-matB-TDH3p) + (tHXT7p-matC-CYC1t) XI-3:: (TEF1p-AtPAL2-FBA1t) X-2::(TEF1-NLS_FapR-ADH1t)* | NAG64 | This work |
| NAG73 | *MATa ura3-52 can1Δ::cas9-natNT2 TRP1 LEU2 HIS3 XII-2:: TDH3p-AtC4H-CYC1t)+(tHXT7p-AtATR2-pYX212t)+(PGK1p-CYB5-ADH1t) X-3::(TPI1p-EcaroL-pYX212t)+(ADH1t-ARO7* $^{G141S}$*-TEF1p)+(PGK1p-ARO4* $^{K229L}$*-CYC1t) X-4::(CYC1t-ARO1-TPI1p)+(TDH3p-ARO2-ADH1t)+(TDH2t-ARO3-TEF1p) XII-4::(TDH3p-At4CL-ADH1t) + (TDH2t-RsCHS-CCW12p)+(tHXT7p-PsCHI-FBAt) XII-1::(TDH3p-At4CL-ADH1t) + (TDH2t-RsCHS-CCW12p)+(tHXT7p-PsCHI-FBAt) XII-5::(pYX212t-PsCHI-PGKp)+ (TEF1p-RsCHS-FBAt) + (CYC1t-At4CL-TPIp) XI-1::(pYX212t-PsCHI-PGKp)+ (TEF1p-RsCHS-FBAt) XI-2::(TPIp-ACC1* $^{S659A,S1157A}$ *-TDH2t) XII-3::(TDH2t-matB-TDH3p) + (tHXT7p-matC-CYC1t) XI-3:: (TEF1_BS123p-AtPAL2-FBA1t) X-2::(TEF1-NLS_FapR-ADH1t)* | NAG65 | This work |
| NAG74 | *MATa ura3-52 can1Δ::cas9-natNT2 TRP1 LEU2 HIS3 XII-2:: TDH3p-AtC4H-CYC1t)+(tHXT7p-AtATR2-pYX212t)+(PGK1p-CYB5-ADH1t) X-3::(TPI1p-EcaroL-pYX212t)+(ADH1t-ARO7* $^{G141S}$*-TEF1p)+(PGK1p-ARO4* $^{K229L}$*-CYC1t) X-4::(CYC1t-ARO1-TPI1p)+(TDH3p-ARO2-ADH1t)+(TDH2t-ARO3-TEF1p) XII-4::(TDH3p-At4CL-ADH1t) + (TDH2t-RsCHS-CCW12p)+(tHXT7p-PsCHI-FBAt) XII-1::(TDH3p-At4CL-ADH1t) + (TDH2t-RsCHS-CCW12p)+(tHXT7p-PsCHI-FBAt) XII-5::(pYX212t-PsCHI-PGKp)+ (TEF1p-RsCHS-FBAt) + (CYC1t-At4CL-TPIp) XI-1::(pYX212t-PsCHI-PGKp)+ (TEF1p-RsCHS-FBAt) XI-2::(TPIp-ACC1* $^{S659A,S1157A}$ *-TDH2t) XII-3::(TDH2t-matB-TDH3p) + (tHXT7p-matC-CYC1t) XI-3:: (TDH3_BS23p-AtPAL2-FBA1t) X-2::(TEF1-NLS_FapR-ADH1t) FDC1::( pYX212t-DCR1-TDH3p) + (tHXT7p-AGO1-CYC1t)* | NAG71 | This work |
| NAG75 | *MATa ura3-52 can1Δ::cas9-natNT2 TRP1 LEU2 HIS3 XII-2:: TDH3p-AtC4H-CYC1t)+(tHXT7p-AtATR2-pYX212t)+(PGK1p-CYB5-ADH1t) X-3::(TPI1p-EcaroL-pYX212t)+(ADH1t-ARO7* $^{G141S}$*-TEF1p)+(PGK1p-ARO4* $^{K229L}$*-CYC1t) X-4::(CYC1t-ARO1-TPI1p)+(TDH3p-ARO2-ADH1t)+(TDH2t-ARO3-TEF1p) XII-4::(TDH3p-At4CL-ADH1t) + (TDH2t-RsCHS-CCW12p)+(tHXT7p-PsCHI-FBAt) XII-1::(TDH3p-At4CL-ADH1t) + (TDH2t-RsCHS-CCW12p)+(tHXT7p-PsCHI-FBAt) XII-5::(pYX212t-PsCHI-PGKp)+ (TEF1p-RsCHS-FBAt) + (CYC1t-At4CL-TPIp) XI-1::(pYX212t-PsCHI-PGKp)+ (TEF1p-RsCHS-FBAt) XI-2::(TPIp-ACC1* $^{S659A,S1157A}$ *-TDH2t) XII-3::(TDH2t-matB-TDH3p) + (tHXT7p-matC-CYC1t) XI-3:: (TDH3_BS23p-AtPAL2-FBA1t) X-2::(TEF1-NLS_FapR-ADH1t) FDC1::( pYX212t-DCR1-TDH3p) + (tHXT7p-AGO1-CYC1t) XI-5::(TDH3_BS23p-FAS1_200bpshRNA-TDH2t)* | NAG74 | This work |
| NAG76 | *MATa ura3-52 can1Δ::cas9-natNT2 TRP1 LEU2 HIS3 XII-2:: TDH3p-AtC4H-CYC1t)+(tHXT7p-AtATR2-pYX212t)+(PGK1p-CYB5-ADH1t) X-3::(TPI1p-EcaroL-pYX212t)+(ADH1t-ARO7* $^{G141S}$*-TEF1p)+(PGK1p-ARO4* $^{K229L}$*-CYC1t) X-4::(CYC1t-ARO1-TPI1p)+(TDH3p-ARO2-ADH1t)+(TDH2t-ARO3-TEF1p) XII-4::(TDH3p-At4CL-ADH1t) + (TDH2t-RsCHS-CCW12p)+(tHXT7p-PsCHI-FBAt) XII-1::(TDH3p-At4CL-ADH1t) + (TDH2t-RsCHS-CCW12p)+(tHXT7p-PsCHI-FBAt) XII-5::(pYX212t-PsCHI-PGKp)+ (TEF1p-RsCHS-FBAt) + (CYC1t-At4CL-TPIp) XI-1::(pYX212t-PsCHI-PGKp)+ (TEF1p-RsCHS-FBAt) XI-2::(TPIp-ACC1* $^{S659A,S1157A}$ *-TDH2t) XII-3::(TDH2t-matB-TDH3p) + (tHXT7p-matC-CYC1t) XI-3:: (TDH3_BS23p-AtPAL2-FBA1t) X-2::(TEF1-NLS_FapR-ADH1t) FDC1::( pYX212t-DCR1-TDH3p) + (tHXT7p-AGO1-CYC1t) XI-5::(TDH3_BS23p-FAS1_250bpshRNA-TDH2t)* | NAG74 | This work |
| IMX581-N4 | *MATa ura3-52 can1Δ::cas9-natNT2 TRP1 LEU2 HIS3 XII-4::(TDH3p-At4CL-ADH1t) + (TDH2t-RsCHS-CCW12p)+(tHXT7p-PsCHI-FBAt)* | IMX581 | This work |

**Supplementary Table 2. Plasmids used in this study.**

| Plasmid ID | Relevant characteristics | Origin |
|---|---|---|
| pCfB1018 | *Template for AtPAL2 and AtC4H* | (3) |
| pCfB0848 | *Template for AtATR2 and CYB5* | (3) |
| pCfB0854 | *Template for At4CL1* | (4) |
| pCfB0826 | *Template for* $ARO4^{K229L}$ *and* $ARO7^{G141S}$ | (3) |
| pCfB2747 | *Template for EcoroL* | (3) |
| pAD | *Template for* $ACC1^{S659A,\ S1157A}$ | (5) |
| pFDA8 | *Template for NLS-FapR* | (6) |
| pX&Y19 | *Template for pCCW12_BS2* | (7) |
| pX&Y22 | *Template for pCCW12_BS123* | (7) |
| pX&Y31 | *Template for pTDH3_BS23* | (7) |
| pFDA7 | *Template for pTEF1_BS123* | (6) |
| pGK3 | *Template for AGO1 and DCR1* | (8) |
| pQC007 | *2µm ampR KlURA3 gRNA-XII-2.Y* | (2) |
| pQC010 | *2µm ampR KlURA3 gRNA-XII-4.Y* | (2) |
| pQC005 | *2µm ampR KlURA3 gRNA-X-3.Y* | (2) |
| pQC008 | *2µm ampR KlURA3 gRNA-X-4.Y* | (2) |
| pQC032 | *2µm ampR URA3 gRNA-XII-1.Y [2x]* | (2) |
| pQC033 | *2µm ampR URA3 gRNA-XII-5.Y [2x]* | (2) |
| pQC030 | *2µm ampR URA3 gRNA-XI-1.Y [2x]* | (2) |
| pQC130 | *2µm ampR URA3 gRNA-XI-2.Y [2x]* | (2) |
| pQC133 | *2µm ampR URA3 gRNA-XII-3.Y [2x]* | (2) |
| pQC029 | *2µm ampR URA3 gRNA-X-2.Y [2x]* | (2) |
| pQC006 | *2µm ampR KlURA3 gRNA-XI-3.Y* | (2) |
| pQC009 | *2µm ampR KlURA3 gRNA-XI-5.Y* | (2) |
| pMEL10 | *2µm ampR KlURA3 gRNA-CAN1.Y* | (2) |
| pMC005 | *2µm ampR URA3 gRNA-AtPAL2.Y* | This work |

**Supplementary Table 3. Primers used in this study.**

| Plasmid ID | Name | Sequence (5'-3') |
|---|---|---|
| **Primers for homologous regions for expression module integration** | | |
| P001 | *XII-4 up-F* | GTATCCGGCTGTTCCTTCATAG |
| P002 | *XII-4 up-R (with TDH3p-F)*[a] | CTTTGAAATGGCAGTATTGATAATGATAAACTCGATGCCATAGTATGTGTGATGG |
| P003 | *XII-4 dn-F (with FBA1t-R)* | CGAGTTCTTTGTAAAGTCTTTCATAGTAGCTTACTATTCCCCATTAGAGTCAAATAAAAG |
| P004 | *XII-4 dn-R* | TTTCTGCCGTACCTGGATGGTCATTTC |
| P005 | *XII-1 up-F* | GTTGAGCTCTGTCCTTCATGG |
| P006 | *XII-1 up-R1 (with TDH3p-F)* | CTTTGAAATGGCAGTATTGATAATGATAAACTCGAGAAAGAACCGAACCGATGCC |
| P007 | *XII-1 dn-F1 (with ADH1t-R)* | AAATCGCTCCCCATTTCACCCAATTGTAGATATGCCTTCCCGTGAATCAACTGCAC |
| P008 | *XII-1 dn-R* | CAATCCTCGCATTTCAGCTTC |
| P009 | *XII-1 up-R2 (with pYX212t-R)* | GCTCCCTTTAGGGTTCCGATTTAGTGGTTTACGGCGAAAGAACCGAACCGATGC |
| P010 | *XII-1 dn-F2 (with FBA1t-R)* | CGAGTTCTTTGTAAAGTCTTTCATAGTAGCTTACTGTTCAGTTTAGTGCTCTGTCTGAG |
| P011 | *XII-5 up-F* | GTAGTGATCATTGGCTTAAC |
| P012 | *XII-5 up-R1 (with pYX212t-R)* | GCTCCCTTTAGGGTTCCGATTTAGTGGTTTACGGCGTGACAATAAATTCAAACCGGT |
| P013 | *XII-5 dn-F1 (with TPI1p-F)* | ATTCTAAGTAAGTTAAATATCCGTAATCTTTAAACCAACTCAGAAGTTTGACAGC |
| P014 | *XII-5 dn-R* | CTCTTTTGCCTTTCAAAAAAG |
| P015 | *XII-5 up-R2 (with CYC1t-R)* | GGACGCTCGAAGGCTTTAATTTGCGGCCGGTACCCGTGACAATAAATTCAAACCGGT |
| P016 | *XII-5 dn-F2 (with FBA1t-R)* | CGAGTTCTTTGTAAAGTCTTTCATAGTAGCTTACTCAACTCAGAAGTTTGACAGC |
| P017 | *XII-5 dn-F3 (with PGK1p-F)* | ATGCCTATTGTGCAGATGTTATAATATCTGTGCGTCAACTCAGAAGTTTGACAGC |
| P018 | *XII-5 up-R3 (with TEF1p-F)* | AGAGTAAAAAAGGAGTAGAAACATTTTGAAGCTATGTGACAATAAATTCAAACCGGT |
| P019 | *XI-1 up-F* | ATTTGTGTGAAGGAATAGTGACG |
| P020 | *XI-1 up-R1 (with pYX212t-R)* | GCTCCCTTTAGGGTTCCGATTTAGTGGTTTACGGCCAATGGGCTTGGTATTCCG |
| P021 | *XI-1 dn-F1 (with TPI1p-F)* | ATTCTAAGTAAGTTAAATATCCGTAATCTTTAAACTTTCTTGGCATTGGCAAATC |
| P022 | *XI-1 dn-R* | AAGAGCCGAGTCCCCATCAG |
| P023 | *XI-1 up-R2 (with CYC1t-R)* | GGACGCTCGAAGGCTTTAATTTGCGGCCGGTACCCCAATGGGCTTGGTATTCCG |
| P024 | *XI-1 dn-F2 (with FBA1t-R)* | CGAGTTCTTTGTAAAGTCTTTCATAGTAGCTTACTTTTCTTGGCATTGGCAAATC |
| P025 | *XI-1 dn-F3 (with PGK1p-F)* | ATGCCTATTGTGCAGATGTTATAATATCTGTGCGTTTTCTTGGCATTGGCAAATC |
| P026 | *XI-1 up-R3 (with TEF1p-F)* | AGAGTAAAAAAGGAGTAGAAACATTTTGAAGCTATCAATGGGCTTGGTATTCCG |
| P027 | *XII-2 up-F* | CGGCATGCAAACATCTACAC |
| P028 | *XII-2 up-R (with GPM1p-F)* | GCTCACAAATCTTAAAGTCATACATTGCACGACTACATAACGCGTTACACGGAAG |
| P029 | *XII-2 dn-F (with PGK1p-F)* | GCAAATGCCTATTGTGCAGATGTTTATAATATCTGTGCGTCTACTATCGGCGACTCTCTC |
| P030 | *XII-2 dn-R* | GAGCGAACGTAAGAGAGGTTAATG |
| P031 | *X-3 up-F* | CGAGATCTTTGTGTTCGGTTACC |
| P032 | *X-3 up-R (with TPI1p-F)* | CTAAGTAAGTTAAATATCCGTAATCTTTAAACGTCTCGTATGTCGGCTCTCGC |
| P033 | *X-3 dn-F (with CYC1t-R)* | GACGCTCGAAGGCTTTAATTTGCGGCCGGTACCCTGTGTCCGCGTTTTCTAAGGC |
| P034 | *X-3 dn-R* | GAGGTGGTTATTGATCACCGGA |
| P035 | *X-4 up-F* | CCCAAAGCTAAGAGTCCCATT |
| P036 | *X-4 up-R (with CYC1t-R)* | GACGCTCGAAGGCTTTAATTTGCGGCCGGTACCCCTGCTCTTGAATGGCGACAG |
| P037 | *X-4 dn-F (with TEF1p-F)* | GAAGAGTAAAAAAGGAGTAGAAACATTTTGAAGCTATAACAGGCATGGGAAGATTCG |
| P038 | *X-4 dn-R* | CTGGTGAGGATTTACGGTATGA |
| P039 | *X-2 up-F* | CGTCTATGAGGAGACTGTTAGTTG |
| P040 | *X-2 up-R (with GPM1p-F)* | GCTCACAAATCTTAAAGTCATACATTGCACGACTAGACCACTTCGAGAGCAAGTTG |
| P041 | *X-2 dn-F (with CYC1t-R)* | GACGCTCGAAGGCTTTAATTTGCGGCCGGTACCCCCTGCATAATCGGCCTCAC |
| P042 | *X-2 dn-R* | CTCGCCAAGGCATTACCATC |
| P043 | *XI-2 up-F* | TAACTCTTCGTATGAGGATTTTC |
| P044 | *XI-2 up-R* | TTCTATGGCACATTTTTCTGTTG |
| P127 | *XI-2 dn-F* | CCACAAGTAAAGCTCGTTGAC |
| P128 | *XI-2 dn-R* | ATGGTTGAAAAGGTTACAGAGG |
| P129 | *XII-3 up-F* | TGTGCCCCTTAAAATTCATATAC |
| P130 | *XII-3 up-R (with TDH2t-R)* | TAAAGCACTTAGTATCACACTAATTGGCTTTTCGCGAATGAGCAGGTACCCCTTA |
| P131 | *XII-3 dn-F (with CYC1t-R)* | GGACGCTCGAAGGCTTTAATTTGCGGCCGGTACCCGCATAGAGCTAATTAGGTTTGAG |

| | | |
|---|---|---|
| P132 | *XII-3 dn-R* | GAACTTACAAGCTGATTTTGGT |
| P133 | *X-2 up-R (with TEF1p-F)* | AGAGTAAAAAAGGAGTAGAAACATTTTGAAGCTATGACCACTTCGAGAGCAAGTTG |
| P134 | *X-2 dn-F (with ADH1t-R)* | AAATCGCTCCCCATTTCACCCAATTGTAGATATGCCCTGCATAATCGGCCTCAC |
| P135 | *XI-3 up-F* | AGTTACTTGCTCTATGCGTTTGC |
| P136 | *XI-3 up-R1 (with GPM1p-F)* | GCTCACAAATCTTAAAGTCATACATTGCACGACTAAATCAGACGCACGCTTGGC |
| P137 | *XI-3 dn-F (with FBA1t-R)* | CGAGTTCTTTGTAAAGTCTTTCATAGTAGCTTACTTTACGTGGATTGAGCCAGCA |
| P138 | *XI-3 dn-R* | TGAGAATCCGGACCAGCAGAT |
| P139 | *XI-3 up-R2 (with CCW12p-F)* | GCCCCTTTTGGACTAACCGTGTGGTTCATGGGTGGAATCAGACGCACGCTTGGC |
| P140 | *XI-3 up-R3 (with TDH3p-F)* | CTTTGAAATGGCAGTATTGATAATGATAAACTCGAAATCAGACGCACGCTTGGC |
| P141 | *XI-3 up-R4 (with TEF1p-F)* | AGAGTAAAAAAGGAGTAGAAACATTTTGAAGCTATAATCAGACGCACGCTTGGC |
| P142 | *FDC1 up-F* | CTGAGCATTTTATTACGTTACTCAAC |
| P143 | *FDC1 up-R (with pYX212t-R)* | CCCTTTAGGGTTCCGATTTAGTGGTTTACGGCTTGAATATATAAATTGACAATTCTTTTG |
| P144 | *FDC1 dn-F (with CYC1t-R)* | GGACGCTCGAAGGCTTTAATTTGCGGCCGGTACCCTTGCCATAGACTTTCTACGG |
| P145 | *FDC1 dn-R* | GTCCATTTATTTTTTATGTGCTGTC |
| P146 | *XI-5 up-F* | GCGGAGAAGTCGTTGATAGC |
| P147 | *XI-5 up-R1 (with TDH3p-F)* | CTTTGAAATGGCAGTATTGATAATGATAAACTCGATGGTGCACGGAGTTTATGG |
| P148 | *XI-5 dn-F (with TDH2t-R)* | TAAAGCACTTAGTATCACACTAATTGGCTTTTCGCTTGTAAAACAGGTATTGGCTGC |
| P149 | *XI-5 dn-R* | GATCATAGATCCGGCACTTAGAG |
| **Primers for amplification of promoters and terminators** | | |
| P045 | *TDH3p-F* | TCGAGTTTATCATTATCAATACTGCC |
| P046 | *TDH3p-R* | CATTTTGTTTGTTTATGTGTGTTTATTCGA |
| P047 | *ADH1t-F* | GCGAATTTCTTATGATTTATGATTTTT |
| P048 | *ADH1t-R* | GCATATCTACAATTGGGTGAAATGG |
| P049 | *TDH2t-F1* | ATTTAACTCCTTAAGTTACTTTAATGATTTAG |
| P050 | *TDH2t-R1 (with ADH1t-R)* | AAATCGCTCCCCATTTCACCCAATTGTAGATATGCGCGAAAAGCCAATTAGTGTGATAC |
| P051 | *CCW12p-F* | CCACCCATGAACCACACGG |
| P052 | *CCW12p-R* | CATTTTGTTTATTGATATAGTGTTTAAGCGAATG |
| P053 | *tHXT7p-F1 (with CCW12p-F)* | GCCCCTTTTGGACTAACCGTGTGGTTCATGGGTGGCTCGTAGGAACAATTTCGGG |
| P054 | *tHXT7p-R* | CATTTTTTGATTAAAATTAAAAAAACTTTTTG |
| P055 | *FBA1t-F* | GTTAATTCAAATTAATTGATATAGTTTTTT |
| P056 | *FBA1t-R* | GATACCGTCGACCTCGAGTC |
| P057 | *GPM1p-F* | TAGTCGTGCAATGTATGACTTTAAGA |
| P058 | *GPM1p-R* | CATTGTTTTTATTGTAATATGTGTGTTTGTT |
| P059 | *CYC1t-F* | GATACCGTCGACCTCGAGTC |
| P060 | *CYC1t-R1* | GGGTACCGGCCGCAAATTAA |
| P061 | *pYX212t-F* | TAGGGCCCACAAGCTTACG |
| P062 | *pYX212t-R1 (with ADH1t-R)* | CAAATCGCTCCCCATTTCACCCAATTGTAGATATGCGCCGTAAACCACTAAATCGGA |
| P063 | *ADH1t-F* | GCGAATTTCTTATGATTTATGATTTTT |
| P064 | *ADH1t-R* | GCATATCTACAATTGGGTGAAATGG |
| P065 | *PGK1p-F* | ACGCACAGATATTATAACATCTGC |
| P066 | *PGK1p-R* | CATTTTGTTATATTTGTTGTAAAAAGTAGATAA |
| P067 | *TPI1p-F* | GTTTAAAGATTACGGATATTTAACTTAC |
| P068 | *TPI1p-R* | CATTTTTAGTTTATGTATGTGTTTTTTGTAG |
| P069 | *TEF1p-F1* | ATAGCTTCAAAATGTTTCTACTCCT |
| P070 | *TEF1p-R* | CATTTTGTAATTAAAACTTAGATTAGATTGC |
| P071 | *TEF1p-F2 (with PGK1p-F)* | ATGCCTATTGTGCAGATGTTATAATATCTGTGCGTATAGCTTCAAAATGTTTCTACTCCT |
| P072 | *PGK1p-F* | ACGCACAGATATTATAACATCTGC |
| P073 | *PGK1p-R* | CATTTTGTTATATTTGTTGTAAAAAGTAGATAA |
| P074 | *pYX212t-R2* | GCCGTAAACCACTAAATCGGA |
| P075 | *TDH2t-R2* | GCGAAAAGCCAATTAGTGTGATAC |
| P076 | *tHXT7p-F2* | CTCGTAGGAACAATTTCGGG |
| P077 | *tHXT7p-F3 (with TDH3p-F)* | CTTTGAAATGGCAGTATTGATAATGATAAACTCGACTCGTAGGAACAATTTCGGG |
| P078 | *CYC1t-R2 (with FBA1t-R)* | TCGAGTTCTTTGTAAAGTCTTTCATAGTAGCTTACGGGTACCGGCCGCAAATTAA |

| | | |
|---|---|---|
| P079 | *TEF1p-F3 (with XI-2_up)* | CGGGGTGTAACTCAACAGAAAAATGTGCCATAGAAATAGCTTCAAAATGTTTCTACTCCT |
| P080 | *TEF1p-R (with mACC1_up)* | GAGAAGACTCGAATAAGCTTTCTTCGCTCATTTTGTAATTAAAACTTAGATTAGATTGC |
| P081 | *TDH2t-F2 (with mACC1_dn)* | AAGAAAAATTGTTGAAGACTTTGAAATAAATTTAACTCCTTAAGTTACTTTAATGATTTAG |
| P082 | *TDH2t-R3 (with XI-2_dn)* | CAACTGATCAACTGGTCAACGAGCTTTACTTGTGGGCGAAAAGCCAATTAGTGTGATAC |
| **Primers for amplification of genes** | | |
| P164 | *At4CL1-F (with TDH3p-R)* | ACTTAGTTTCGAATAAACACACATAAACAAACAAAATGGCTCCACAAGAACAAG |
| P165 | *At4CL1-R (with ADH1t-F)* | TTAATAATAAAAATCATAAATCATAAGAAATTCGCTCACAAACCGTTAGCCAAC |
| P166 | *RsCHS-F (with CCW12p-R)* | CTGTCATTCGCTTAAACACTATATCAATAAACAAAATGGTTACTGTTGAAGATGTTAG |
| P167 | *RsCHS-R (with TDH2t-F)* | AAACTAAATCATTAAAGTAACTTAAGGAGTTAAATTTAAGTACACAATGAATGCAAAAC |
| P168 | *PsCHI-F (with tHXT7p-R)* | CAAAAACAAAAAGTTTTTTTAATTTTAATCAAAAAATGGCAAAACCACCATCTG |
| P169 | *PsCHI-R (with FBA1t-F)* | TCATTAAAAAACTATATCAATTAATTTGAATTAACTTATTTTAACAATTCAGAAATTCTTGC |
| P170 | *HaCHS-F (with CCW12p-R)* | CTGTCATTCGCTTAAACACTATATCAATAAACAAAATGGTTACTGTTGAAGAAG |
| P171 | *HaCHS-R (with TDH2t-F)* | AAACTAAATCATTAAAGTAACTTAAGGAGTTAAATTTAATTAATAGCAACTGAATG |
| P172 | *SmCHS-F (with CCW12p-R)* | CTGTCATTCGCTTAAACACTATATCAATAAACAAAATGGTTACTGTTGAAGAATAC |
| P173 | *SmCHS-R (with TDH2t-F)* | AAACTAAATCATTAAAGTAACTTAAGGAGTTAAATTTAAGCTGCAACAGAATGC |
| P174 | *PhCHI-F (with tHXT7p-R)* | CAAAAACAAAAAGTTTTTTTAATTTTAATCAAAAAATGTCTCCACCAGTTTCAG |
| P175 | *PsCHI-R (with FBA1t-F)* | TCATTAAAAAACTATATCAATTAATTTGAATTAACTTAAACACCAATAACTGGG |
| P176 | *SmCHI-F (with tHXT7p-R)* | CAAAAACAAAAAGTTTTTTTAATTTTAATCAAAAAATGGCTGCTGTTACTAAATTG |
| P177 | *SmCHI-R (with FBA1t-F)* | TCATTAAAAAACTATATCAATTAATTTGAATTAACATGGCTGCTGTTACTAAATTG |
| P178 | *AtPAL2-F (with GPM1p-R)* | CCAAACAAACACACATATTACAATAAAAACAATGGATCAAATCGAAGCTATGTTG |
| P179 | *AtPAL2-R (with FBA1t-F)* | CTCATTAAAAAACTATATCAATTAATTTGAATTAACTCAGCAGATAGGAATAGGAGCACC |
| P180 | *AtC4H-F (with TDH3p-R)* | GTTTCGAATAAACACACATAAACAAACAAAATGGACTTGTTGTTGTTGGAAAAG |
| P181 | *AtC4H-R (with CYC1t-F)* | CATAACTAATTACATGACTCGAGGTCGACGGTATCTCAACAGTTTCTTGGCTTCATAAC |
| P182 | *AtATR2-F (with tHXT7p-R)* | CAAAAACAAAAAGTTTTTTTAATTTTAATCAAAAAATGTCCTCCTCTTCTTCATCATC |
| P183 | *AtATR2-R (with pYX212t-F)* | CCCGGGTCGACGCGTAAGCTTGTGGGCCCTATCACCAGACATCTCTCAAGTATCTAC |
| P184 | *CYB5-F (with PGK1p-R)* | GTAATTATCTACTTTTTACAACAAATATAACAAAATGCCTAAAGTTTACAGTTACCAAG |
| P185 | *CYB5-R (with ADH1t-F)* | TTTAATAATAAAAATCATAAATCATAAGAAATTCGCTCATTCGTTCAACAAATAATAAGC |
| P186 | *EcoroL-F (with TPI1p-R)* | CTATAACTACAAAAAACACATACATAAACTAAAAATGACACAACCTCTTTTTCTGATC |
| P187 | *EcoroL-R (with pYX212t-F)* | GATACCCGGGTCGACGCGTAAGCTTGTGGGCCCTATCAACAATTGATCGTCTGTGCC |
| P188 | *ARO7*-F (with TEF1p-R)* | CATAGCAATCTAATCTAAGTTTTAATTACAAAATGGATTTCACAAAACCAGA |
| P189 | *ARO7*-R (with ADH1t-F)* | TATTTAATAATAAAAATCATAAATCATAAGAAATTCGCTTACTCTTCCAACCTTCTTAGC |
| P190 | *ARO4*-F (with PGK1p-R)* | GTAATTATCTACTTTTTACAACAAATATAACAAAATGAGTGAATCTCCAATGTTCG |
| P191 | *ARO4*-R (with CYC1t-F)* | CATAACTAATTACATGACTCGAGGTCGACGGTATCCTATTTCTTGTTAACTTCTCTTCTT |
| P192 | *ARO1-F (with TPI1p-R)* | CTATAACTACAAAAAACACATACATAAACTAAAAATGGTGCAGTTAGCCAAAGTC |
| P193 | *ARO1-R (with CYC1t-F)* | CATAACTAATTACATGACTCGAGGTCGACGGTATCCTACTCTTTCGTAACGGCATCA |
| P194 | *ARO2-F (with TDH3p-R)* | GTTTCGAATAAACACACATAAACAAACAAAATGTCAACGTTTGGGAAACTG |
| P195 | *ARO2-R (with ADH1t-F)* | CTTATTTAATAATAAAAATCATAAATCATAAGAAATTCGCTTAATGAACCACGGATCTGGA |
| P196 | *ARO3-F (with TEF1p-R)* | CATAGCAATCTAATCTAAGTTTTAATTACAAAATGTTCATTAAAAACGATCACGC |
| P197 | *ARO3-R (with TDH2t-F)* | CTAAATCATTAAAGTAACTTAAGGAGTTAAATCTATTTTTTCAAGGCCTTTCTTC |
| P198 | *PHA2-F (with GPM1p-R)* | CCAAACAAACACACATATTACAATAAAAACAATGGCCAGCAAGACTTTGAGG |
| P199 | *PHA2-R (with CYC1t-F)* | ACTAATTACATGACTCGAGGTCGACGGTATCTTATTTGTGATAATATCTCTCATTTCTG |
| P200 | *At4CL1-F (with TPI1p-R)* | TCTATAACTACAAAAAACACATACATAAACTAAAAATGGCTCCACAAGAACAAG |
| P201 | *At4CL1-R (with CYC1t-F)* | CATAACTAATTACATGACTCGAGGTCGACGGTATCTCACAAACCGTTAGCCAAC |
| P202 | *RsCHS-F (with TEF1p-R)* | AAGCATAGCAATCTAATCTAAGTTTTAATTACAAAATGGTTACTGTTGAAGATGTTAG |
| P203 | *RsCHS-R (with FBA1t-F)* | TCATTAAAAAACTATATCAATTAATTTGAATTAACTTAAGTACACAATGAATGCAAAAC |
| P204 | *PsCHI-F (with PGK1p-R)* | AGTAATTATCTACTTTTTACAACAAATATAACAAAATGGCAAAACCACCATCTG |
| P205 | *PsCHI-R (with pYX212t-F)* | ACCCGGGTCGACGCGTAAGCTTGTGGGCCCTATTATTTTAACAATTCAGAAATTCTTG |
| P206 | *mACC1_up 500bp-F* | ATGAGCGAAGAAAGCTTATTCG |
| P207 | *mACC1_up 500bp-R* | CAATAGTGGATTCTCGGAGGC |
| P208 | *mACC1_dn 500bp-F* | ATCGTGAGAGAGAACTATTGCC |
| P209 | *mACC1_dn 500bp-R* | TTATTTCAAAGTCTTCAACAATTTTTCTTTATC |
| P210 | *matB-F (with TDH3p-R)* | CTTAGTTTCGAATAAACACACATAAACAAACAAAATGTCTAATCATTTGTTTGATGCAATG |
| P211 | *matB-R (with TDH2t-F)* | AAACTAAATCATTAAAGTAACTTAAGGAGTTAAATTTAAGTTCTTGTGTACAAATCAGC |

| | | |
|---|---|---|
| P212 | *matC-F (with tHXT7p-R)* | CAAAAACAAAAAGTTTTTTTAATTTTAATCAAAAAATGGGTATCGAATTGTTGTCTATC |
| P213 | *matC-R (with CYC1t-F)* | TAACTAATTACATGACTCGAGGTCGACGGTATCTTAAACTAAACCTGGAACAACAAAAAC |
| P223 | *AtPAL2-F (with CCW12p-R)* | CTGTCATTCGCTTAAACACTATATCAATAAACAAAATGGATCAAATCGAAGCTATG |
| P224 | *AtPAL2-F (with TEF1p-R)* | TAGCAATCTAATCTAAGTTTTAATTACAAAATGGATCAAATCGAAGCTATG |
| P225 | *AtPAL2-F (with TDH3p-R)* | TTTCGAATAAACACACATAAACAAACAAAATGGATCAAATCGAAGCTATG |
| P226 | *NLS_FapR-F (with TEF1p-R)* | GCAATCTAATCTAAGTTTTAATTACAAAATGCCAAAGAAGAAGAGAAAGGTTAG |
| P227 | *NLS_FapR-R (with ADH1t-F)* | TTAATAATAAAAATCATAAATCATAAGAAATTCGCTTATGAATGTTTTGAACGATACATG |
| P228 | *DCR1-F (with TDH3p-R)* | GATACCCGGGTCGACGCGTAAGCTTGTGGGCCCTATCACAGATTGTTGCAATGCC |
| P229 | *DCR1-R (with pYX212t-F)* | ACTTAGTTTCGAATAAACACACATAAACAAACAAAATGAATAGAGAAAAAAGCGCC |
| P230 | *AGO1-F (with tHXT7p-R)* | CAAAAACAAAAAGTTTTTTTAATTTTAATCAAAAAATGTCATCCAATTCGGAGGAG |
| P231 | *AGO1-R (with CYC1t-F)* | CATAACTAATTACATGACTCGAGGTCGACGGTATCTCATATGTAGTACATGATGTCAG |
| P232 | *FAS1_sense 200bp-F (with TDH3p-R)* | TTTCGAATAAACACACATAAACAAACAAAATGGACGCTTACTCCACAAG |
| P233 | *FAS1_ sense 200bp-R (with rad9_intron1-F)* | GAAAAAAGTTCCAACACACCTGCTGCAGCAAACCCTTCAG |
| P234 | *rad9_intron1-F* | CAGGTGTGTTGGAACTTTTTTCAAACCTTACTAAACATTGAAACTAATTGGTAAAGATA |
| P235 | *rad9_intron1-R* | TATCTTTACCAATTAGTTTCAATGTTTAGTAAGGTTTGAAAAAAGTTCCAACACACCTG |
| P236 | *FAS1_antisense 200bp-F (with rad9_intron1-R)* | AACATTGAAACTAATTGGTAAAGATACTGCAGCAAACCCTTCAG |
| P237 | *FAS1_antisense 200bp-R (with TDH2t-F)* | AAACTAAATCATTAAAGTAACTTAAGGAGTTAAATATGGACGCTTACTCCACAAG |
| P238 | *FAS1_ sense 250bp-F (with TDH3p-R)* | TTTCGAATAAACACACATAAACAAACAAAATGGACGCTTACTCCACAAG |
| P239 | *FAS1_ sense 250bp-R (with rad9_intron1-F)* | GAAAAAAGTTCCAACACACCTGAGACCTGATCGAATTGACC |
| P240 | *FAS1_antisense 250bp-F (with rad9_intron1-R)* | AACATTGAAACTAATTGGTAAAGATAAGACCTGATCGAATTGACC |
| P241 | *FAS1_antisense 250bp-R (with TDH2t-F)* | AAACTAAATCATTAAAGTAACTTAAGGAGTTAAATATGGACGCTTACTCCACAAG |
| **Primers for *AtPAL2* gene deletion and repair oligos** | | |
| P214 | *Vector backbone-F* | GATCATTTATCTTTCACTGCGGAGAAG |
| P215 | *Vector backbone-R* | GTTTTAGAGCTAGAAATAGCAAGTTAAAATAAGGCTAGTC |
| P216 | *Sequencing-F* | CACCTTTCGAGAGGACGATG |
| P217 | *Sequencing-R* | GCTGGCCTTTTGCTCACATG |
| P218 | *AtPAL2_gRNA* | TGCGCATGTTTCGGCGTTCGAAACTTCTCCGCAGTGAAAGATAAATGATCAATGGTACAGCTGTTGGTTCGTTTTAGAGCTAGAAATAGCAAGTTAAAATAAGGCTAGTCCGTTATCAAC |
| P219 | *Repair Fragment 1-F* | GCCATTTTTTTTTCTGTATCGGGCCCTCCTTACTGCTCTCCTTCCGTGTAACGCGTTATGAATTTTTATTTCATTCTGGAACTCTTCGAGTTCTTTGTAAAGTCTTTCATAGTAGCTTAC |
| P220 | *Repair Fragment 1-R* | GTAAGCTACTATGAAAGACTTTACAAAGAACTCGAAGAGTTCCAGAATGAAATAAAAATTCATAACGCGTTACACGGAAGGAGAGCAGTAAGGAGGGCCCGATACAGAAAAAAAAATGGC |
| P221 | *Repair Fragment 2-F* | GCCATTTTTTTTTCTGTATCGGGCCCTCCTTACTGCTCTCCTTCCGTGTAACGCGTTATGCTACTATCGGCGACTCTCTCGAAATTTTTCTTAACGCGTCCTTGTACTGCGTCTAACGCT |
| P222 | *Repair Fragment 2-R* | AGCGTTAGACGCAGTACAAGGACGCGTTAAGAAAAATTTCGAGAGAGTCGCCGATAGTAGCATAACGCGTTACACGGAAGGAGAGCAGTAAGGAGGGCCCGATACAGAAAAAAAAATGGC |

[a] short overlap to the fragments in the parentheses were included in corresponding primers.

**Supplementary Table 4. Codon optimized genes used in this study.**

| Gene | Sequence (5'-3') |
|---|---|
| *HaCHS* | ATGGTTACTGTTGAAGAAGTTAGAAAAGCACAAAGAGCTGAAGGTCCAGCAACTGTTATGGCTATTGGTACAGCAGTT<br>CCACCAAATTGTGTTGATCAAGCTACTTACCCAGATTACTACTTCAGAATTACAAATTCTGAACATAAGGCAGAATTAA<br>AGGAAAAGTTCCAAAGAATGTGTGATAAATCACAAATTAAGAAAAGATATATGTACTTGAATGAAGAAGTTTTAAAAGA<br>AAATCCAAATATGTGTGCATACATGGCTCCATCTTTAGATGCTAGACAAGATATCGTTGTTGTTGAAGTTCCAAAATTG<br>GGTAAAGAAGCTGCTGTTAAAGCAATTAAAGAATGGGGTCAACCAAAGTCTAAGATCACTCATTTGGTTTTCTGTACT<br>ACATCAGGTGTTGATATGCCAGGTGCTGATTACCAATTGACAAAATTGTTGGGTTTAAGACCATCAGTTAAGAGATTG<br>ATGATGTACCAACAAGGTTGTTTTGCAGGTGGTACAGTTTTGAGATTGGCTAAAGATTTGGCAGAAAACAATAAGGGT<br>GCTAGAGTTTTGGTTGTTTGTTCTGAAATCACTGCTGTTACTTTTAGAGGTCCAACTGATACACATTTGGATTCATTAG<br>TTGGTCAAGCATTGTTTGGTGACGGTGCTGCTGCTATTATTATTGGTTCTGATCCAATTCCAGAAGTTGAAAAGCCATT<br>GTTCGAATTGGTTTCTGCTGCTCAAACTATTTTACCAGATTCAGAAGGTGCTATTGATGGTCATTTGAGAGAAGTTGGT<br>TTAACATTCCATTTGTTGAAGGATGTTCCAGGTTTGATCTCTAAAAATGTTGAAAAATCATTGACTGAAGCCTTTAAAC<br>CATTGGGTATCTCTGATTGGAACTCTTTATTTTGGATTGCTCATCCAGGTGGTCCAGCAATTTTGGATCAAGTTGAAG<br>CTAAATTGTCTTTAAAGCCAGAAAAATTGAGAGCAACAAGACATGTTTTGTCAGAATACGGTAACATGTCTTCAGCTTG<br>TGTTTTGTTTATTTTAGATGAAATGAGAAGAAAGTCTAAGGAAGATGGTTTGAAGACTACAGGTGAAGGTATTGAATG<br>GGGTGTTTTGTTTGGTTTTGGTCCAGGTTTAACTGTTGAAACAGTTGTTTTGCATTCAGTTGCTATTAATTAA |
| *RsCHS* | ATGGTTACTGTTGAAGATGTTAGAAGAGCACAAAGAGCTGAAGGTCCAGCAACAGTTATGGCTATTGGTACTGCAAC<br>ACCATCTAATTGTGTTGATCAATCAACTTACCCAGATTTCTACTTTAGAATTACAAATTCTGAACATAAGGCTGAATTAA<br>AGGAAAAGTTCCAAAGAATGTGTGATAAATCAATGATTAAGAAAAGATATATGTACTTAACTGAAGAAATTTTGAAGGA<br>AAACCCATCTGTTTGTGAATATATGGCTCCATCATTAGATGCAAGACAAGATATGGTTGTTGTTGAAGTTCCAAAATTG<br>GGTAAAGAAGCTGCAACTAAAGCTATTAAAGAATGGGGTCAACCAAAGTCTAAGATCACACATTTGGTTTTCTGTACT<br>ACATCAGGTGTTGATATGCCAGGTGCAGATTACCAATTGACTAAATTGTTGGGTTTAAGACCATCTGTTAAGAGATTG<br>ATGATGTACCAACAAGGTTGTTTTGCTGGTGGTACAGTTTTGAGATTGGCTAAAGATTTGGCAGAAAACAATAAGGGT<br>GCTAGAGTTTTGGTTGTTTGTTCAGAAATCACTGCTGTTACTTTTAGAGGTCCATCTGATACACATTTGGATTCATTGG<br>TTGGTCAAGCATTGTTTGGTGACGGTGCTGCTGCTATTATTGTTGGTGCAGATCCAGTTCCAGAAGTTGAAAAGCCAT<br>TGTTCGAATTGGTTTCTGCTGCTCAAACTATTTTACCAGATTCAGATGGTGCTATTGATGGTCATTTGAGAGAAGTTGG<br>TTTGACATTCCATTTGTTGAAGGATGTTCCAGGTTTGATCTCTAAAAATATCGAAAAAGCATTGACTGAAGCATTTCAA<br>CCATTGGGTATTTCTGATTGGAACTCTATTTTCTGGATTGCTCATCCAGGTGGTCCAGCAATTTTGGATCAAGTTGAAT<br>TGAAATTATCTTTGAAGCCAGAAAAATTGAGAGCTACAAGACATGTTTTGTCAGAATACGGTAACATGTCTTCAGCATG<br>TGTTTTGTTTATTTTGGATGAAATGAGAAGAAAATCTGCTGAAGAAGGTTTAAAAACTACAGGTGAAGGTTTAGAATGG<br>GGTGTTTTGTTTGGTTTTGGTCCAGGTTTGACTGTTGAAACAGTTGTTTTGCATTCATTGTGTACTTAA |
| *SmCHS* | ATGGTTACTGTTGAAGAATACAGAAAGGCACAAAGAGCTGAAGGTCCAGCAACAGTTATGGCTATTGGTACTTCTACA<br>CCATCAAATTGTGTTGATCAATCTGCTTACCCAGATTATTACTTTAGAATTACTAAATCAGAACATAAGACAGAATTAAA<br>GGAAAAGTTTAAAAGAATGTGTGAAAAATCTATGATTAAGAAAAGATATATGTACTTAACTGAAGAAATTTTGAAAGAA<br>AAATCTAATATGTGTGCATATATGGCTCCATCATTAGATGCAAGACAAGATATCGTTGTTGTTGAAGTTCCAAAATTGG<br>GTAAAGAAGCTGCACAAAAAGCTATTAAAGAATGGGGTCAACCAAAGTCTAAGATCACTCATTTGGTTTTCTGTACTA<br>CATCAGGTGTTGATATGCCAGGTTGTGATTACCAATTGACAAAATTGTTGGGTTTAAGACCATCTGTTAAGAGATTGAT<br>GATGTACCAACAAGGTTGTTTTGCTGGTGGTACAGTTTTGAGATTGGCAAAAGATTTGGCTGAAAACAATAAGGGTGC<br>AAGAGTTTTGGTTGTTTGTTCTGAAATCACTGCTGTTACTTTTAGAGGTCCATCTGAATCACATTTGGATTCATTGGTT<br>GGTCAAGCATTGTTTGGTGACGGTGCTGCTGCTGTTATTATTGGTTCTGATCCAATTTCAGGTGTTGAAAGACCATTA<br>TTTGAATTGGTTTCTGCTGCTCAAACTTTGTTACCAGATTCAGAAGGTGCAATTGATGGTCATTTGAGAGAAGTTGGTT<br>TGACATTCCATTTGTTGAAAGATGTTCCAGGTATCATCTCTAAAAATATCGATAAATCATTAGTTGAAGCATTTCAACCA<br>TTGGGTATCTCTGATTGGAACTCTTTATTTTGGATTGCACATCCAGGTGGTCCAGCTATTTTGGATCAAGTTGAATTGA<br>AATTGGGTTTGAACCCAGAAAAATTGAGAGCAACTAGAGAAGTTTTGTCTAACTACGGTAACATGTCTTCAGCTTGTG<br>TTTTGTTTATTTTGGATGAAATGAGAAAAGCATCAGCTAAACAAGGTTTACCAACTACAGGTGAAGGTTTACAATGGG<br>GTGTTTTGTTTGGTTTTGGTCCAGGTTTGACTGTTGAAACATTAGTTTTGCATTCTGTTGCAGCTTAA |
| *PhCHI* | ATGTCTCCACCAGTTTCAGTTACTAAGATGCAAGTTGAAAACTATGCTTTTGCACCAACAGTTAATCCAGCAGGTTCTA<br>CTAATACATTGTTTTTAGCTGGTGCAGGTCATAGAGGTTTGGAAATCGAGGGTAAATTCGTTAAGTTTACTGCTATCG<br>GTGTTTACTTGGAAGAATCTGCTATCCCATTTTTAGCAGAAAAGTGGAAGGGTAAAACTCCACAAGAATTAACAGATT<br>CAGTTGAATTTTTCAGAGATGTTGTTACTGGTCCATTCGAAAAGTTTACTAGAGTTACAATGATCTTGCCATTGACAGG<br>TAAACAATACTCTGAAAAGGTTGCTGAAAATTGTGTTGCACATTGGAAAGGTATTGGTACTTACACAGATGATGAAGG<br>TAGAGCTATCGAAAAGTTTTTGGATGTTTTTAGATCAGAAACATTTCCACCAGGTGCATCTATCATGTTCACTCAATCA<br>CCATTAGGTTTGTTGACAATTTCTTTTGCTAAAGATGATTCAGTTACTGGTACAGCTAACGCAGTTATTGAAAATAAGC<br>AATTGTCTGAAGCTGTTTTAGAATCAATCATCGGTAAACATGGTGTTTCTCCAGCTGCAAAATGTTCAGTTGCTGAAAG<br>AGTTGCAGAATTGTTGAAGAAATCTTATGCTGAAGAAGCATCAGTTTTCGGTAAACCAGAAACTGAAAAATCTACAATC<br>CCAGTTATTGGTGTTTAA |
| *PsCHI* | ATGGCAAAACCACCATCTGTTTCAGGTGTTAACATCGAATCTTATGCTTTTCCACCAACTGTTAAACCACCAGGTTCTA<br>CTAAAACATTGTTTTTAGGTGGTGCAGGTGTTAGAGGTTTGGAAGTTCCAACTGGTCAATTCGTTAAGTTTACTGCTAT<br>CGGTGTTTACTTGGAAGATAACGCAATCACTTCATTAGCTGTTAAGTGGAAAGGTAAAACTGCTGAAGAATTAACAGA<br>ATCTGATGATTTCTTTAGAGATATCGTTACAGGTCCATTCGAAAAGTTTACTCAAGTTACAATGATCTTGCCATTGACT |

| | |
|---|---|
| | GGTCAACAATACTCAGAAAAGGTTACAGAAAACTGTGTTGCTTACTGGAAAGCAGTTGGTGCTTACACTGATGCTGAA<br>GCATCTGCTATTGAAAAGTTTATTGAAGTTTTTAAAGATGAAAAGTTTCCACCAGGTTCTTCAATTTTGTTTACTCAAAC<br>ACCAGAAGGTTCTTTAACAATCGGTTTTTCAAAGGATGGTGTTTTGCCAGAAGTTGGTAATGCAGTTGTTGAAAATAA<br>GCAATTGTCTGAAGCTGTTTTAGAATCAATCATCGGTAAACATGGTGTTTCTCCAGAAGCTAAACAATCATTAGCTGCA<br>AGAATTTCTGAATTGTTAAAATAA |
| *SmCHI* | ATGGCTGCTGTTACTAAATTGCAAATCGAAAACTACGTTTTTCCACCAACTATGGTTAAGCCAGTTGGTTCTAACAACA<br>CTTTCTTTTTGGCTGGTGCAGGTTCAAGAGGTTTAGAAATCGAAGGTAGATTCGTTAAGTTTACTGCTATCGCAGTTTA<br>CTTGGAAGAATCTGCTATTCCATTTTTAGCTGCAAAGTGGAAGGGTAAATCTTCAGAAGAATTAACTGATTCAGTTGAA<br>TTTTTCAAGGATATCGTTACAGGTCCATTCGAAAAGTTTACTCAAGTTACAATGATCTTGCCATTGACTGGTAAACAAT<br>ACTCTGAAAAGGTTGCTGAAAACTGTGTTGCTAACTGGAAAGCAATTGGTACTTACTCTGATGCTGAATCACAAGCAA<br>TCGAAAAGTTCTTGAACGTTTTCCAATCTGAAACATTCCCACATGGTGCATCAATTTTGTTTACTCAATCTCCATTGGG<br>TTCATTAACAATTTCTTTTTCAAAGGATGATTCTGTTCCATCAATCGGTAACGCTGTTATTGAAAATAAGCAATTGTCTG<br>AAGCAGTTTTAGATTCAATCATCGGTAAACATGGTGTTTCTCCAGCTGCAAAGTGTTCAATCGCTAAGAGAGTTTCTG<br>AATTGTTAGAAAAATCAAATGCTGAAGCAGTTGTTAATGAAAAACCAGGTTCTGGTTTGTCAACAATTCAATAA |
| *RtmatB* | ATGTCTAATCATTTGTTTGATGCAATGAGAGCTGCAGCTCCTGGTAATGCTCCTTTTATTAGAATCGATAACACTAGAA<br>CATGGACTTACGATGATGCATTTGCTTTATCTGGTAGAATTGCATCAGCTATGGATGCATTGGGTATTAGACCAGGTG<br>ACAGAGTTGCTGTTCAAGTTGAAAAATCTGCAGAAGCTTTGATCTTGTATTTGGCATGTTTGAGATCAGGTGCTGTTTA<br>TTTGCCATTGAATACAGCATACACTTTGGCTGAATTGGATTACTTCATCGGTGACGCAGAACCAAGATTGGTTGTTGT<br>TGCTTCTTCAGCAAGAGCTGGTGTTGAAACTATTGCAAAACCAAGAGGTGCTATTGTTGAAACATTAGATGCAGCTGG<br>TTCTGGTTCATTGTTAGATTTGGCAAGAGATGAACCAGCTGATTTTGTTGATGCATCTAGATCAGCTGATGATTTGGCA<br>GCTATTTTGTACACTTCTGGTACTACAGGTAGATCAAAAGGTGCAATGTTGACACATGGTAATTTGTTGTCAAACGCTT<br>TGACTTTGAGAGATTTTTGGAGAGTTACAGCAGGTGACAGATTGATCCATGCTTTGCCAATCTTCCATACTCATGGTTT<br>ATTCGTTGCTACAAACGTTACTTTGTTAGCAGGTGCTTCTATGTTTTTGTTGTCAAAGTTCGATCCAGAAGAAATCTTG<br>TCTTTGATGCCACAAGCTACTATGTTGATGGGTGTTCCAACATTCTACGTTAGATTGTTGCAATCACCAAGATTGGATA<br>AGCAAGCAGTTGCTAACATCAGATTGTTTATTTCTGGTTCAGCACCATTGTTAGCTGAAACACATACTGAATTTCAAGC<br>AAGAACTGGTCATGCTATTTTAGAAAGATACGGTATGACAGAAACTAACATGAACACTTCTAACCCATACGAAGGTAA<br>AAGAATTGCTGGTACAGTTGGTTTTCCATTGCCAGATGTTACAGTTAGAGTTACTGATCCAGCAACAGGTTTAGCTTT<br>GCCACCAGAACAAACTGGTATGATCGAAATTAAAGGTCCAAACGTTTTTAAAGGTTACTGGAGAATGCCAGAAAAGAC<br>TGCAGCTGAATTCACTGCTGATGGTTTCTTTATTTCTGGTGACTTGGGTAAAATCGATAGAGATGGTTACGTTCATATT<br>GTTGGTCGTGGTAAAGATTTGGTTATTTCTGGTGGTTACAACATCTATCCAAAGGAAGTTGAAGGTGAAATCGATCAA<br>ATCGAAGGTGTTGTTGAATCAGCTGTTATTGGTGTTCCACATCCAGATTTTGGTGAAGGTGTTACTGCTGTTGTTGTTA<br>GAAAACCAGGTGCAGCTTTGGATGAAAAGGCAATCGTTTCTGCTTTGCAAGATAGATTGGCAAGATACAAGCAACCA<br>AAGAGAATCATCTTCGCTGAAGATTTGCCAAGAAATACAATGGGTAAAGTTCAAAAGAATATCTTGAGACAACAATATG<br>CTGATTTGTACACAAGAACTTAA |
| *RtmatC* | ATGGGTATCGAATTGTTGTCTATCGGTTTGTTGATTGCTATGTTCATCATCGCAACTATCCAACCAATTAATATGGGTG<br>CTTTGGCATTTGCTGGTGCATTTGTTTTAGGTTCTATGATCATCGGTATGAAGACAAACGAAATCTTCGCTGGTTTCCC<br>ATCAGATTTGTTTTTGACTTTGGTTGCAGTTACATATTTGTTCGCTATCGCACAAATTAATGGTACTATCGATTGGTTG<br>GTTGAATGTGCTGTTAGATTAGTTAGAGGTAGAATTGGTTTGATTCCATGGGTTATGTTCTTGGTTGCTGCAATCATCA<br>CAGGTTTCGGTGCTTTGGGTCCAGCTGCAGTTGCTATTTTGGCACCAGTTGCTTTGTCTTTCGCAGTTCAATACAGAA<br>TCCATCCAGTTATGATGGGTTTAATGGTTATTCATGGTGCTCAAGCAGGTGGTTTTTCTCCAATCTCAATCTATGGTGG<br>TATCACTAACCAAATCGTTGCTAAAGCAGGTTTACCATTTGCTCCAACATCTTTGTTTTTATCTTCATTTTTCTTTAATTT<br>GGCTATCGCAGTTTTGGTTTTCTTTGTTTTCGGTGGTGCTAGAGTTATGAAACATGATCCAGCATCATTGGGTCCATTA<br>CCAGAATTGCATCCAGAAGGTGTTTCTGCTTCAATTAGAGGTCATGGTGGTACTCCAGCTAAACCAATTAGAGAACAT<br>GCATACGGTACTGCTGCAGATACAGCTACTACATTGAGATTGAACAACGAAAGAATCACTACATTGATCGGTTTGACA<br>GCTTTAGGTATCGGTGCATTGGTTTTTAAGTTTAATGTTGGTTTAGTTGCAATGACTGTTGCTGTTGTTTTGGCATTGT<br>TATCTCCAAAGACACAAAAGGCTGCAATCGATAAAGTTTCTTGGTCAACTGTTTTGTTGATCGCTGGTATCATCACATA<br>CGTTGGTGTTATGGAAAAAGCTGGTACTGTTGATTACGTTGCAAATGGTATTTCTTCATTGGGTATGCCATTGTTGGTT<br>GCTTTGTTGTTGTGTTTCACTGGTGCAATTGTTTCAGCTTTTGCATCTTCAACAGCTTTATTGGGTGCAATCATCCCAT<br>TGGCTGTTCCATTTTTATTGCAAGGTCATATCTCTGCTATTGGTGTTGTTGCTGCAATCGCAATCTCAACTACAATCGT<br>TGATACTTCTCCATTTTCAACAAATGGTGCTTTAGTTGTTGCTAATGCACCAGATGATTCTAGAGAACAAGTTTTGAGA<br>CAATTGTTGATCTATTCAGCATTGATCGCTATCATCGGTCCAATTGTTGCTTGGTTGGTTTTTGTTGTTCCAGGTTTAG<br>TTTAA |

**Supplementary Table 5. Expression modules in the integration constructs used in this study**

| ID | DNA fragment[a] |
|---|---|
| M1 | *XII-4 up-TDH3p-**At4CL1**-ADH1t +TDH2t* |
| M2 | *ADH1t +TDH2t-**HaCHS**-CCW12p+tHXT7p-**PhCHI**-FBAt- XII-4 dn* |
| M3 | *ADH1t +TDH2t-**HaCHS**-CCW12p+tHXT7p-**PsCHI**-FBAt- XII-4 dn* |
| M4 | *ADH1t +TDH2t-**HaCHS**-CCW12p+tHXT7p-**SmCHI**-FBAt- XII-4 dn* |
| M5 | *ADH1t +TDH2t-**RsCHS**-CCW12p+tHXT7p-**PhCHI**-FBAt- XII-4 dn* |
| M6 | *ADH1t +TDH2t-**RsCHS**-CCW12p+tHXT7p-**PsCHI**-FBAt- XII-4 dn* |
| M7 | *ADH1t +TDH2t-**RsCHS**-CCW12p+tHXT7p-**SmCHI**-FBAt- XII-4 dn* |
| M8 | *ADH1t +TDH2t-**SmCHS**-CCW12p+tHXT7p-**PhCHI**-FBAt- XII-4 dn* |
| M9 | *ADH1t +TDH2t-**SmCHS**-CCW12p+tHXT7p-**PsCHI**-FBAt- XII-4 dn* |
| M10 | *ADH1t +TDH2t-**SmCHS**-CCW12p+tHXT7p-**SmCHI**-FBAt- XII-4 dn* |
| M11 | *X-3 up-TPI1p-**EcaroL**-pYX212t-ADH1t-**ARO7** $^{G141S}$-TEF1p-PGK1p-**ARO4** $^{K229L}$-CYC1t- X-3 dn* |
| M12 | *X-4 up-CYC1t-**ARO1**-TPI1p-TDH3p-**ARO2**-ADH1t-TDH2t-**ARO3**-TEF1p-X-4 dn* |
| M13 | *X-2 up-GPM1p-**PHA2**-CYC1t-X-2 dn* |
| M14 | *XII-1 up-TDH3p-**At4CL1**-ADH1t- XII-1 dn* |
| M15 | *XII-1 up-TDH2t-**RsCHS**-CCW12p-tHXT7p-**PsCHI**-FBAt-XII-1 dn* |
| M16 | *XII-1 up-TDH3p-**At4CL1**-ADH1t-TDH2t* |
| M17 | *ADH1t-TDH2t-**RsCHS**-CCW12p-tHXT7p-**PsCHI**-FBAt-XII-1 dn* |
| M18 | *XII-5 up-pYX212t-**PsCHI**-PGKp-TEF1p-**RsCHS**-FBAt-CYC1t* |
| M19 | *FBAt-CYC1t-**At4CL1**-TPIp-XII-5 dn* |
| M20 | *XII-5 up-CYC1t-**At4CL1**-TPIp-XII-5 dn* |
| M21 | *XII-5 up-TEF1p-**RsCHS**-FBAt-XII-5 dn* |
| M22 | *XII-5 up-pYX212t-**PsCHI**-PGKp-XII-5 dn* |
| M23 | *XII-5 up-pYX212t-**PsCHI**-PGKp-TEF1p-**RsCHS**-FBAt-XII-5 dn* |
| M24 | *XI-1 up-pYX212t-**PsCHI**-PGKp-TEF1p-**RsCHS**-FBAt-CYC1t* |
| M25 | *FBAt-CYC1t-**At4CL1**-TPIp-XI-1 dn* |
| M26 | *XI-1 up-CYC1t-**At4CL1**-TPIp-XI-1 dn* |
| M27 | *XI-1 up-TEF1p-**RsCHS**-FBAt-XI-1 dn* |
| M28 | *XI-1 up-pYX212t-**PsCHI**-PGKp-XI-1 dn* |
| M29 | *XI-1 up-pYX212t-**PsCHI**-PGKp-TEF1p-**RsCHS**-FBAt-XI-1 dn* |
| M30 | *XI-3 up-TDH2t-**RsCHS**-CCW12p-tHXT7p-**PsCHI**-FBAt-XI-3 dn* |
| M31 | *XI-2 up-TPIp-ACC1 $^{S659A,S1157A}$ up 500bp* |
| M32 | ***ACC1** $^{S659A,S1157A}$* |
| M33 | *ACC1 $^{S659A,S1157A}$ dn 500bp-TDH2t-XI-2 dn* |
| M34 | *XII-3 up-TDH2t-**matB**-TDH3p-tHXT7p* |
| M35 | *TDH3p-tHXT7p-**matC**-CYC1t- XII-3 dn* |
| M36 | *X-2 up-TEF1-**NLS_FapR**-ADH1t-X-2 dn* |
| M37 | *XI-3 up-GPM1p-**AtPAL2**-FBA1t-XI-3 dn* |
| M38 | *XI-3 up-CCW12p-**AtPAL2**-FBA1t-XI-3 dn* |
| M39 | *XI-3 up-CCW12p_BS2-**AtPAL2**-FBA1t-XI-3 dn* |
| M40 | *XI-3 up- CCW12p_BS123-**AtPAL2**-FBA1t-XI-3 dn* |
| M41 | *XI-3 up-TDH3p-**AtPAL2**-FBA1t-XI-3 dn* |
| M42 | *XI-3 up-TDH3p_BS23-**AtPAL2**-FBA1t-XI-3 dn* |
| M43 | *XI-3 up-TEF1p-**AtPAL2**-FBA1t-XI-3 dn* |
| M44 | *XI-3 up-TEF1p_BS123-**AtPAL2**-FBA1t-XI-3 dn* |
| M45 | *FDC1 up-pYX212t-**DCR1**-TDH3p-tHXT7p* |
| M46 | *TDH3p-tHXT7p-**AGO1**-CYC1t- FDC1 dn* |
| M47 | *XI-5 up-TDH3p-FAS1_sense 200bp-rad9_intron1 80bp* |
| M48 | *XI-5 up-TDH3p-FAS1_sense 250bp-rad9_intron1 80bp* |

| | |
|---|---|
| M49 | *XI-5 up-TDH3_BS23p-FAS1_sense 200bp-rad9_intron1 80bp* |
| M50 | *XI-5 up-TDH3_BS23p-FAS1_sense 250bp-rad9_intron1 80bp* |
| M51 | *rad9_intron1 80bp-FAS1_antisense 200bp-TDH2t- XI-5 dn* |
| M52 | *rad9_intron1 80bp-FAS1_antisense 250bp-TDH2t- XI-5 dn* |

[a] Bold font indicates genes expressed; *p*, indicates promoter; *t*, indicates terminator; underline, indicates up-stream (up) and down-stream (dn) sequences for Cas9-mediated homologous recombination.
